# Functional connectivity in infants’ visual cortex: Links to motion processing and autism

**DOI:** 10.1101/2025.06.11.658898

**Authors:** Irzam Hardiansyah, Giorgia Bussu, Sven Bölte, Emily J.H. Jones, Terje Falck-Ytter

## Abstract

In a previously published study, we found atypical visual cortical laterality patterns during global motion perception in 5-month-old infants who showed high levels of autistic symptoms in toddlerhood. Here, using data from a separate experiment, we examined whether these results could reflect altered visual cortical functional connectivity in theta, alpha, and gamma rhythms. We assessed this in a sample of 5- month- old infants (n = 59; 39 elevated familial likelihood of autism) by means of electroencephalography (EEG) when they were watching videos showing social and non-social scenes. Gamma connectivity between midline and far-lateral visual cortex when viewing social scenes was linked to both later autism symptoms and global motion visual cortical laterality we reported in the previous study. This may indicate a shared integrative mechanism underlying social perception and global motion processing. Further, we found that higher midline-to-lateral theta connectivity in the visual cortex when perceiving non-social scenes in infancy was strongly associated with having more autistic symptoms at follow up, but uncorrelated with concurrent motion perception. Our study points to atypical functional connectivity in the visual cortex as a potential early marker of autistic symptoms and highlights a probable link between motion processing and social perception.

## INTRODUCTION

Although it is well established that the emergence of autistic symptoms is linked to differences in brain development, much remains to be understood concerning the specific nature of these differences. Both altered structural and functional brain development have been implicated^1-4^, and one infant brain imaging study suggested that the emergence of autistic symptoms was linked to a hyper-expansion of the cortex across different areas in the brain^5^, including the visual cortex. Consistently, we recently found atypical activation of the visual cortex during low-level global perceptual processing^6^ in infants with high levels of autistic symptoms in toddlerhood^7^. This study highlighted the potential developmental implication of atypical motion processing in the visual cortex of autistic children. These findings from infants are also in line with a recent study of transcriptomic alterations in the visual cortex of autistic adults^8^ (see also ref^9^).

Altered connectivity has been postulated to be a functional brain signature in autism^10-13^, and it is possible that the above-mentioned differences in global motion processing are associated with differences in connectivity in the visual cortex among infants later diagnosed with autism. Therefore, we investigated the functional connectivity in the visual cortex in a cohort of 5-month-old infants from the Early Autism in Sweden (EASE). The cohort resembled the one from earlier study^7^, but was larger, and included additional participants that had reached the study endpoint (see **Results**, also **Supplementary Tables 1-2** for analysis-specific exclusions and sample sizes). We aimed to assess i) if atypical functional connectivity in the visual cortex is linked to later autism diagnosis and ii) if there is a correlation between the connectivity and global motion processing. Specifically, we performed analyses of within-visual cortex resting-state connectivity based on infants’ EEG data, using the debiased Weighted Phase-Lag Index (dbWPLI)^14^, a phase-lag measure that had been reported to be robust for use with infants^15,16^. An analysis plan was registered prior to data analysis https://osf.io/e8y5x.

Given that our previous findings indicated atypical balance between midline and lateral (peripheral) activity in the visual cortex during motion processing in infants subsequently showing autistic symptoms^7^, we focussed on patterns of synchronization between the primary visual cortex (V1) and lateral extrastriate cortices (**Fig. 1c**). We assessed within-visual cortex EEG connectivity by adapting a graph-based approach that has previously been used with fMRI data^17,18^. Our approach is similar to that used in ref^19^ but with simplified graph measures instead of the traditional ones (e.g., path length, clustering, etc.). These simplified measures were defined in our analysis plan and allowed us to make specific inferences about communications between midline and lateral areas of the visual cortex (see full descriptions in **Methods**).

**Figure 1.**
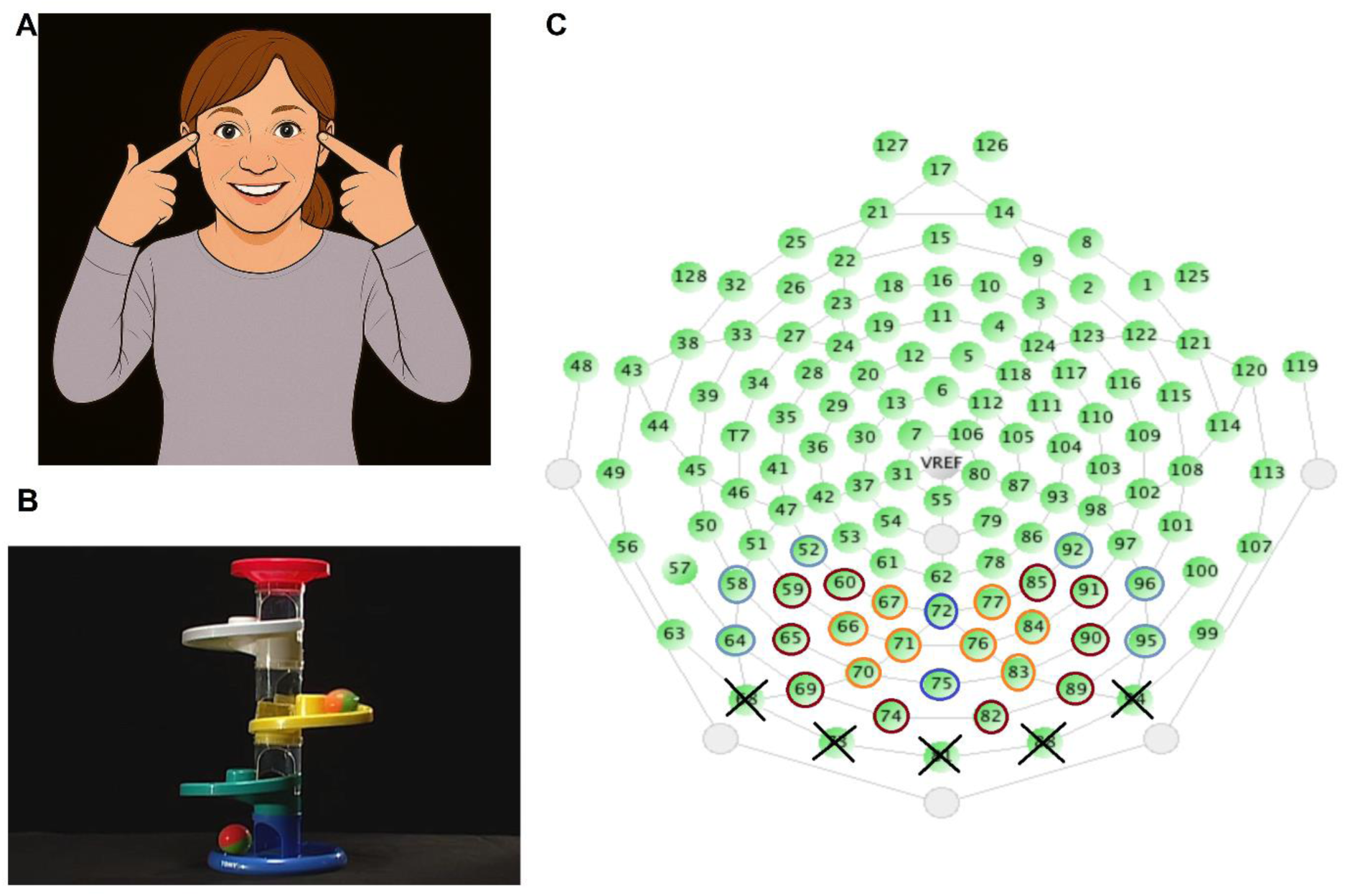
Experimental setup. **a.** A replacement figure for a screenshot of stimulus video clip of the social condition, generated using an A.I. image generator based on the original photo (done due to restriction on bioRxiv to put photos on published manuscript). **b.** Screenshots from the non-social (toy) stimulus video clip. **c.** EEG electrode layout and the part of occipital scalp region that we defined as visual cortex (described fully in **Methods**) were divided into 6 areas of interest (AoIs) relative to, but excluding, the two midline “seed” electrodes coloured in blue. These 6 AoIs extended bilaterally and covered successively further regions of the occipital cortex, forming step-1 (yellow), step-2 (green), and step-3 (purple) block of electrodes, where the “step” reflected the average (binned) Euclidian distance of the corresponding block from the seed electrodes. Crossed channels were discarded. The Euclidean distances were calculated from the 3-dimensional coordinates of each electrode estimated by EEGLab. A mean dbWPLI functional connectivity score was then calculated for each of the 6 AoIs relative to the seed region, and scores for the step-1 area (left and right, separately) were defined as the Near-Connectivity and those for the step-3 area (left and right, separately) as the Far-Connectivity. *Note:* The photo is from an actor / research assistant who has consented to the usage of her image in this article.

The infants were watching short movie clips, either showing a social (a woman singing nursery rhymes; **Fig. 1a**) or a non-social (colourful rotating toys; **Fig. 1b**) scene. Thus, in contrast to our previous experimental stimuli used to probe global motion processing specifically^7^, the current stimuli showed visual motion in a (for infants) ecologically meaningful context. We assessed functional connectivity of theta, alpha, and gamma brain rhythms; see **Methods** for more details. The three frequency bands were slightly shifted to adjust for the young age of the participants and were chosen due to their being the most commonly investigated brain rhythms in the contexts of cognitive development^20,21^, social attention^22^, and autism^23-25^. Furthermore, infants’ theta and alpha rhythms has long been investigated^26,27^ and shown to possess similar functional significance as the analogous rhythms in adults^28,29^. We chose not to assess functional connectivity during the previously reported global motion experiment^7^ because of the strong oscillations elicited by the stimuli themselves (alternating between signal and noise several times per second).

As in the previous report^7^, we compared three groups: i) infants with elevated likelihood of autism and significant later autistic symptoms (EL-High ADOS), ii) infants with elevated autism likelihood and few or no later symptoms (EL-Low ADOS), iii) and infants with low likelihood of autism (LL). The categorization of autistic symptom presentation within the EL group was based on established cut offs on the infants’ Calibrated Severity Scores (CSS)^30^ from the Autism Diagnostic Observation Schedule 2^nd^ edition (ADOS-2)^31^ assessed at 36 months of age (see **Methods**). Following our analysis plan, we created analytical variables called the Near-Connectivity and Far-Connectivity, defined as the score of dbWPLI connectivity from the midline occipital cortex to the lateral occipital region immediately adjacent to it (“step-1 distance”) and from the midline to the furthest lateral occipital region (“step-3 distance”) included in our areas of interest, respectively (see **Fig. 1c**, also described fully in **Methods**). Thus, in analogy to the previous Laterality Score^7^, the two variables should enable us to probe whether atypical balance exists between functional connectivity in the proximity of V1 and that in the lateral (extrastriate) areas. Finally, we used the same Global Motion Laterality Score variable as in the previous work^7^ to represent (lateral) global motion processing and analyzed its relation to the dbWPLI connectivity variables we derived in this study. (Notably, in this paper we use the term “laterality” to denote peripheral versus central (midline) area of the visual cortex, rather than left versus right.)

## RESULTS

### Sample sizes and descriptive statistics

Due to the different nature of EEG data pre-processing and analysis to compute the Near-/Far-Connectivity and Global Motion Laterality Score^7^ variables, the sample size differed for the analyses: N_Connectivity_ = 68, N_LatScore_ = 95 (**Supplementary Tables 1-2**). Descriptive statistics for each resulting dataset are presented in **Supplementary Tables 3-4**.

### Topographical distribution of midline-to-lateral functional connectivity within the visual cortex

Plots of topographical functional connectivity across the 6 AoIs defined in **Fig. 1c** are shown in **Fig. 2** (*n* = 59; *n*_EL-HiADOS_ = 11, *n*_EL-LoADOS_ = 28, *n*_LL_ = 20). The left column (**2a, c, e**) showed individual topographical plots, while the right column (**2b, d, f**) show group average topographies. The plots are presented in the chronology of our analysis steps (see **Methods**): original dbWPLI scores (top panel, **2a, b**), vector-normalized dbWPLI scores (middle panel, **2c, d**), flipped (all maxima arbitrarily allocated to the right side) and normalized dbWPLI scores (bottom panel, **2e, f**).

**Figure 2.**
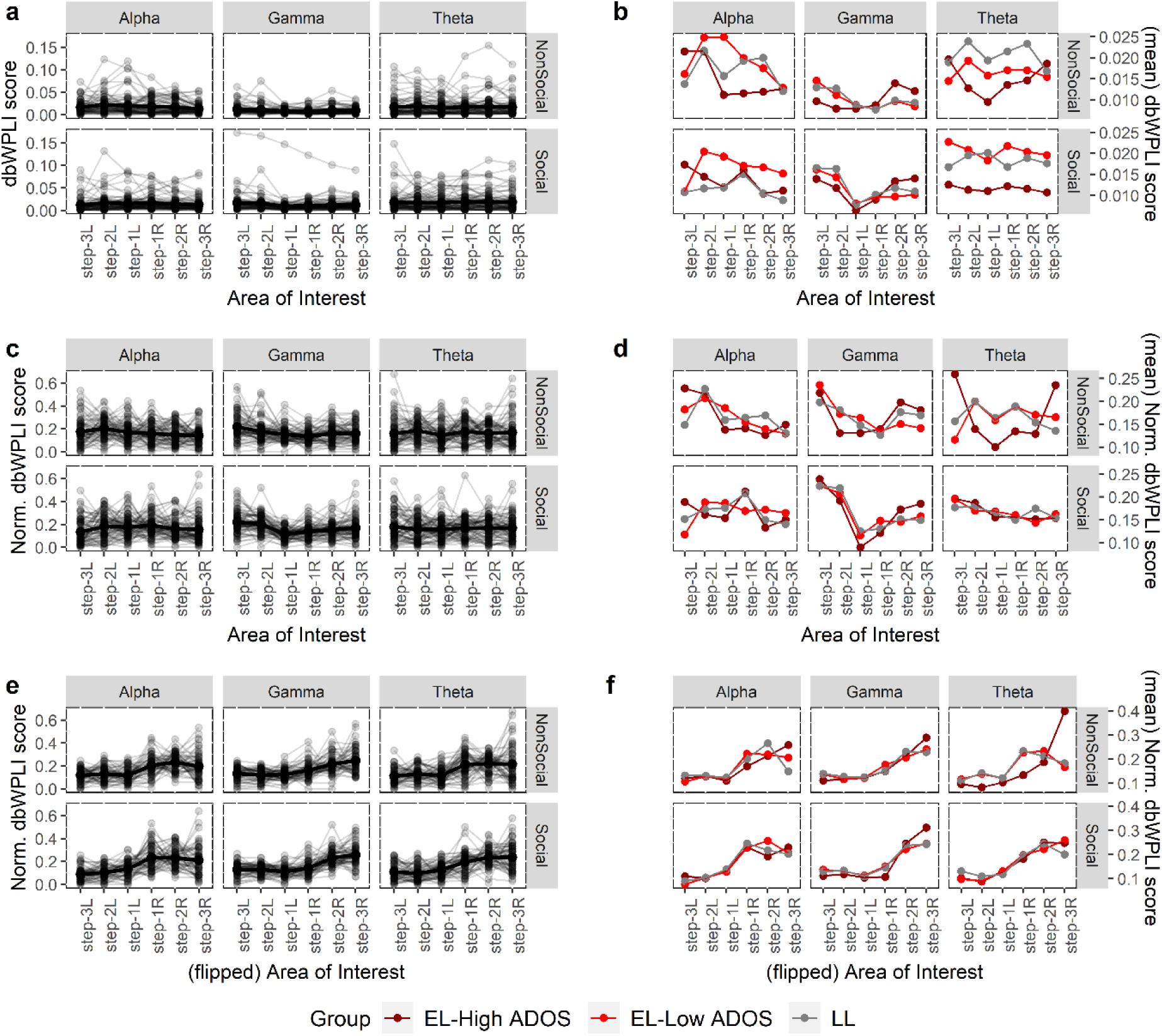
Topographical theta, alpha, and gamma connectivity during observation of social and non-social stimuli. **a.** Raw (unnormalized, unflipped) topographical dbWPLI scores across the occipital electrodes in theta, alpha, and gamma rhythms for the two stimulus conditions from all individual participants irrespective of group. **b.** Group average plots (EL-High ADOS = red; EL-Low ADOS = orange; LL = green) from the topographical plots in **a** for the six experimental (3 bands x 2 stimuli) conditions. **c-d.** Corresponding normalized (see **Methods**) topographical dbWPLI scores across the occipital scalp. **e-f.** Corresponding flipped topographical dbWPLI scores. Here all individual maxima are always to the right (hence ‘left’ and ‘right’ are arbitrary; see **Methods**). The bilaterality patterns seen in **b** and **d**, especially for Gamma-NonSocial, Gamma-Social, and Theta-NonSocial conditions, were due to the almost equal numbers of infants having a peak either on the left or right side of the topography, but very rarely on both, which was made apparent in **f**. For all plots, *n* = 59. See Fig. 3c and **Supplementary Fig. 2** for the error bars (for **b,d,f**) and their corresponding non-parametric difference testing for the flipped normalized topographies in **c**.

The EL-High ADOS group had substantially lower overall raw Theta-Social connectivity (lower-right panel of **Fig. 2b**), and a different raw Theta-NonSocial topographic morphology (upper-right panel of **Fig. 2b**) compared to the other two groups. The flipped and normalized plots revealed clear patterns of unilateral, i.e., only right or left but not both, connectivity between midline and lateral occipital areas. The Theta-NonSocial topography for the EL-High ADOS group remained distinctive in the flipped plots (upper-right panel of **Fig. 2f**). See **Supplementary Table 5** for formal tests of across-group differences.

### Association between infants’ visual cortical functional connectivity and autistic symptoms in toddlerhood

For infants who had complete data for dbWPLI scores and ADOS-2 score (*n* = 59; *n*_EL-HiADOS_ = 11, *n*_EL-LoADOS_ = 28, *n*_LL_ = 20), the non-parametric variable selection (see **Methods**) provided 5 connectivity variables –Theta-NonSocial, Gamma-Social, Gamma-NonSocial Far-Connectivities and Theta-NonSocial, Alpha-Social Near-Connectivities– as the most predictive of ADOS-2 CS total score at 36 months within the repeated 10-fold cross-validation scheme. In the subsequent model-fitting (see “*Modelling association between functional connectivity and autism*” in **Methods**), however, only the three Far-Connectivity variables Theta-NonSocial (*β* = 1.12, partial-*η*^2^ = .23, \*\*\**p* < .001, *n* = 59), Gamma-Social (*β* = .60, partial-*η*^2^ = .09, \**p* = .036, *n* = 59), and Gamma-NonSocial (*β* = -.59, partial-*η*^2^ = .08, \**p* = .047, *n* = 59) were found significant, together accounting for ∼44% of variability in ADOS-2 score. Two covariates, autism likelihood status (EL = 1, LL = 0) (*β* = .98, partial-*η*^2^ = .09, \**p* = .034, *n* = 59) and sex (M = 1, F = 0) (*β* = .87, partial-*η*^2^ = .08, \**p* = .047, *n* = 59) were found significant. Following the preregistered analysis plan, we did an analysis of moderation by autism likelihood status. This analysis revealed a significant interaction between likelihood status and Far-Connectivity in the Theta-NonSocial condition, such that the effect is stronger in the EL group (*β* = 1.91, partial-*η*^2^ = .14, \*\**p* = .008, *n* = 59), while in both Gamma-Social and Gamma-NonSocial (both *p* > .070) the effects were no longer significant at α = .050. The resulting model now explained more variability (*R*^2^ = 49%) in ADOS-2 scores compared the main-effect model. See **Supplementary Table 5** for more details on these modelling results.

A follow up analysis revealed that the association of Theta-NonSocial Far-Connectivity with ADOS-2 score was driven by the EL sub-sample (**Fig. 3a**). Further group-level analyses showed that the EL-High ADOS group displayed greater Theta-NonSocial Far-Connectivity scores overall (**Fig. 3b**), which reflect their distinctive average topographical connectivity pattern in that condition (**Fig. 3c**; see also **Supplementary Table 5)**.

**Figure 3.**
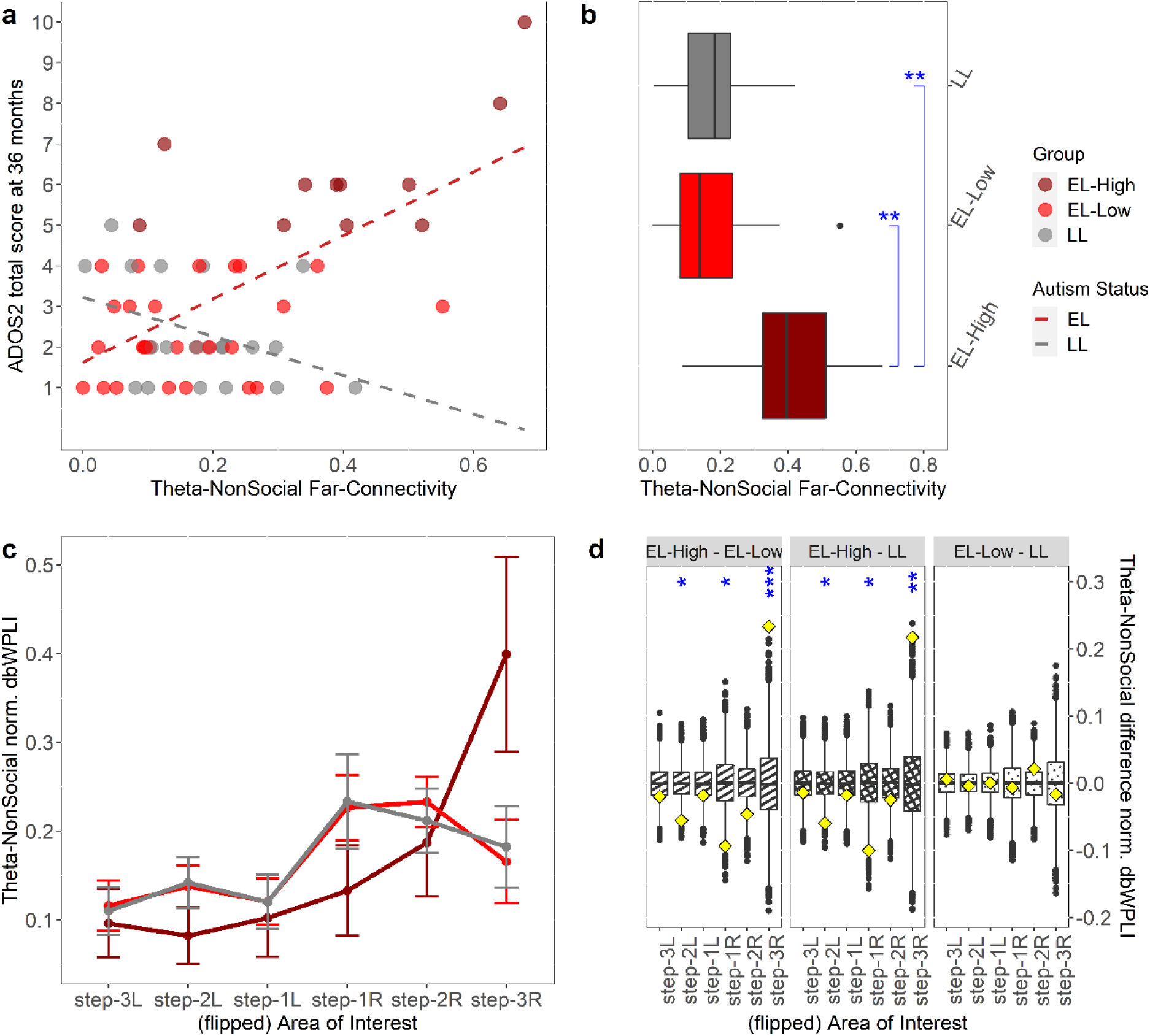
Different midline-to-far lateral visual cortical theta connectivity during non-social stimulus viewing is associated with having more symptoms at three years of age. **a.** Correlation between Theta-NonSocial Far-Connectivity and ADOS-2 CSS total score (*r* = .50, \*\*\**p* < .001, *n* = 59; regression line not shown) was driven by a strong positive correlation (*r* = .64, *\*\*\*p* < .001, *n* = 39) in the EL sub-sample (LL sample: *r* = -.38, *p* = .095, *n* = 20). **b.** Accordingly, group difference analysis of Theta-NonSocial Far-Connectivity revealed significant differences between the EL-High ADOS group (*n* = 11) and both EL-Low ADOS (*d* = 1.61, \*\**p* = .005, *n* = 28) and LL (*d* = 1.57, \*\**p* = .008, *n* = 20) groups, but no difference was seen between the latter two groups (*p* = .492); see also **Supplementary Fig. 1** for plots of the other connectivity variables.. **c.** Mean flipped normalized full topographic patterns of the functional connectivity for the three groups (crf. Fig. 2f), showing that the EL-High ADOS group’s connectivity result is distinctive from the other two groups (see **d** for statistical group comparisons). **d.** Results of non-parametric permutation testing of pairwise group differences (see **Methods**) of topographic patterns shown in **c**, done for each of the six areas of interest making up the full topography, where each panel presents a comparison of two groups (boxplots are simulated distributions using 10,000 randomization of the group labels, diamonds actual point values in our data, stars mark significance); see also **Supplementary Fig. 2** for plots of the other 5 conditions. Compared to EL-Low ADOS infants, the EL-High ADOS infants had on average stronger connectivity in step-3R AoI (*M* = .234, *d* = 4.16, \*\*\**p* < .001) but weaker in step-1R AoI (*M* = -.094, *d* = -2.41, \**p* = .050) and in step-2L AoI (*M* = -.056 *d* = -2.36, \**p* = .050) in the theta band during non-social video viewing. Comparison of EL-High ADOS group with LL infants produced similarly pronounced differences (step-3R AoI: *M* = .217, *d* = 3.67, \*\**p* = .004; step-1R AoI: *M* = -.100, *d* = -2.44, \**p* = .050; step-2L AoI: *M* = -.060, *d* = -2.44, \**p* = .050) in theta band in response to this non-social stimulus. Comparison between EL-Low ADOS and LL infants did not produce any difference (all *p* > .450). All comparisons were FDR-corrected. (Notes: EL = Elevated Likelihood; LL = Low Likelihood; EL-High = EL-High ADOS; EL-Low = EL-Low ADOS; step-*d*L/-R = step-*d*Left/-Right, *d* = 1, 2, 3; * = *p* < .05; ** = *p* < .01; *** = *p* < .001)

### Linking laterality of visual cortical functional connectivity and lateral global motion processing in infancy to later autism

With *n* = 43 (*n*_EL-HiADOS_ = 7, *n*_EL-LoADOS_ = 21, *n*_LL_ = 15) infants who had complete data for Near- and Far-Connectivity, Global Motion Laterality Score, and ADOS-2 score, only Theta-NonSocial (*β* = .56, partial-*η*^2^ = .11, \**p* = .039, *n* = 43) and Gamma-NonSocial (*β* = -.59, partial-*η*^2^ = .11, \**p* = .046, *n* = 43) Far-Connectivities were significant when we refitted the main-effect model of functional connectivity association with autism. The model had a *R*^2^ of 32.2%. We did not fit the interaction model due to low power to detect an interaction effect. Expanding the main-effect model by adding Global Motion Laterality Score as an extra predictor (see “*Inferring relationship between functional connectivity and global motion processing*” in **Methods**), both Theta-NonSocial (*β* = .63, partial-*η*^2^ = .15, \**p* = .017, *n* = 43) and Gamma-NonSocial (*β* = -.57, partial-*η*^2^ = .11, \**p* = .042, *n* = 43) Far-Connectivities were still significant, along with the Global Motion Laterality Score (*β* = .47, partial-*η*^2^ = .15, \**p* = .045, *n* = 43) itself. The model’s *R*^2^ was 39.7% and a subsequent nested likelihood ratio test of the models before and after the expansion showed that this increase in variance explained was significant (χ_1_^2^ = 5.01, *p* = .025). No covariate turned out significant in either model. Notably, comparison of ANOVA results of these two models indicated that variance explained by Global Motion Laterality Score was distinct from that explained by either Theta-NonSocial or Gamma-NonSocial Far-Connectivity; however, it reduced the variance explained attributed to Gamma-Social Far-Connectivity (see **Table 1**). A subsequent analysis of association between Global Motion Laterality Score and all 12 Near-/Far-Connectivity variables, using the same variable selection procedure as in the previous section (also see **Methods**), revealed Gamma-Social Far-Connectivity (*β* = .36, partial-*η*^2^ = .12, \**p* = .021, *n* = 49) as the only connectivity variable significantly associated with Global Motion Laterality Score (**Fig. 4a**), consistent with the above results. Complete modelling results can be found in **Supplementary Table 5**.

**Figure 4.**
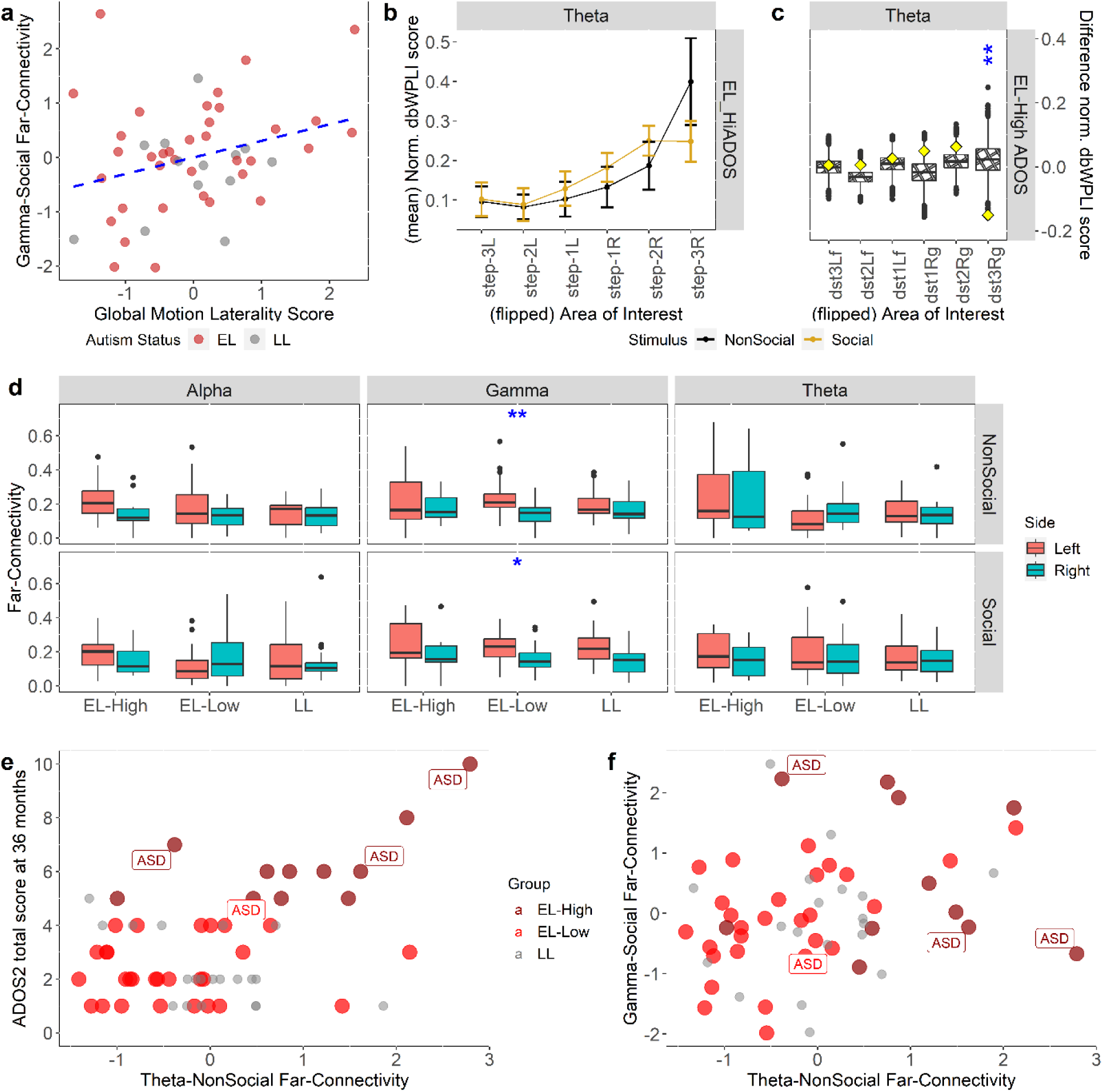
Linking functional connectivity laterality to global motion processing and ASD diagnosis. **a.** Association between Gamma-Social Far-Connectivity and Global Motion Laterality Score was significant (*r* = .30, \**p* = .038, *n* = 49) for all participants who had data for both variables. **b.** Mean flipped normalized topographic patterns of theta functional connectivity for the EL-High ADOS group, showing an elevated far-connectivity during the viewing of non-social relative to social stimuli (see **c** for the formal statistical comparison). **c.** Accordingly, non-parametric permutation testing of between-stimulus difference (see **Methods**) of topographic patterns shown in **c** revealed a significant difference (social – non-social) in Step-3R AoI (*d* = -3.05, \*\**p* =.002, *n* = 11); boxplots are simulated distributions using 10,000 randomization of the stimulus labels, diamonds actual point values in our data, stars mark significance). See also **Supplementary Fig. 3** for the complete plots and permutation testing of between-stimulus differences for all the groups and bands. **d.** Wilcoxon tests of group differences in left or right (i.e., right minus left) connectivity asymmetry was performed to see whether left-sidedness or right-sidedness in any of the three frequency bands and two stimulus types is more related to autism. No group difference was found, except for in gamma frequency band and in both stimulus types (Gamma-NonSocial: *d* = -1.03, \*\**p* = .004, *n* = 58; Gamma-Social: *d* = -.82, \**p* = .031, *n* = 58), but only for the EL-Low ADOS group. In both cases, left asymmetry was more prevalent. Boxplots show the sample median, and the first and third quartiles; whiskers show minimum and maximum (± 1.5 s.d.); dots are outliers (± 1.96 s.d.). **e.** Reproduction of Fig. 3a but now with autism diagnosis cases shown. **f.** Individuals diagnosed with autism are shown as a function of both non-social theta connectivity laterality (x-axis) and social gamma connectivity laterality (y-axis). All cases in EL-High ADOS group (dark red) correspond to extreme laterality in either non-social theta or social gamma connectivity. See **Supplementary Table 5** for numerical details of all results presented here. All *p*-values were FDR-corrected. (Notes: EL = Elevated Likelihood; LL = Low Likelihood; EL-High = EL-High ADOS; EL-Low = EL-Low ADOS; * = *p* < .05; ** = *p* < .01; *** = *p* < .001).

**Table 1.** Analysis of Variance of Functional Connectivity Association Models of ADOS CSS at 36 months without and with Global Motion Laterality Score (GMLS)

| Predictor | Model without GMLS |  | Model with GMLS |  |
| --- | --- | --- | --- | --- |
|  | Sum of Square | *p*-value | Sum of Square | *p*-value |
| Sex | 3.44 | .210 | 1.52 | .381 |
| Autism Likelihood Status | 1.96 | .341 | .82 | .519 |
| Age (5 months) | 7.01 | .077 | 6.68 | .072 |
| Theta-NonSocial Far-Connectivity | 9.67 | .039 | 12.16 | .017 |
| Gamma-NonSocial Far-Connectivity | 9.07 | .046 | 8.60 | .042 |
| Gamma-Social Far-Connectivity | 4.46 | .155 | 1.01 | .474 |
| Global Motion Laterality Score |  |  | 8.37 | .045 |
| Residuals | 76.05 |  | 67.68 |  |
| Degrees of freedom | 36 |  | 35 |  |

As a supplementary analysis, we confirmed that for the expanded sample of *n* = 82 (*n*_EL-HiADOS_ = 19, *n*_EL-LoADOS_ = 44, *n*_LL_ = 19) infants, the previously reported association between Global Motion Laterality Score and ADOS-2 total score at 36 months still held (*β* = .53, partial-*η*^2^ = .07, \*\**p* = .018, *n* = 82); see **Supplementary Table 5**.

### Functional connectivity differences between stimulus types and between left and right hemispheres

Although not specified in the preregistered Analysis Plan, we performed two additional analyses to aid the interpretation of the results: i) social vs. non-social topographic connectivity (both raw and flipped-normalized) differences in each frequency band by group; ii) left/right far-lateral connectivity difference in each experimental condition by group. Only in the flipped topographies, we found lower social relative to non-social theta connectivity at step-3R AoI in EL-High ADOS group (**Figs. 4b, c**), but the reverse (higher social relative to non-social theta connectivity) at the same AoI in the EL-Low ADOS group (**Supplementary Fig. 3**). No lateralization (left vs right) difference was found in any condition or group (**Fig. 4d**). See **Supplementary Table 5** for detailed results.

### Autism diagnosis and non-social theta and social gamma far-lateral connectivity

There were only four individuals diagnosed with autism in our sample, therefore we did not perform a formal analysis of autism diagnosis. Nonetheless, we provided plots (**Figs. 4e, f**) which showed that all but one case corresponds to an extreme score in either Theta-NonSocial or Gamma-Social Far-Connectivity variable.

## DISCUSSION

We investigated infants’ visual cortical functional connectivity and tested its link to both our earlier global motion processing findings^7^ and to later presentation of autistic symptoms. As the current study was motivated by our previous report^7^ on global motion processing in the visual cortex, we focussed exclusively on this area of the brain. We found that a stronger functional connectivity between midline and far lateral areas of the visual cortex in gamma band during viewing of social scenes (smiling and singing women) was associated with a stronger discrepancy of lateral-to-midline activity during global motion processing, as well as with later presentation of autistic symptoms. Our results also revealed that infants’ midline-to-far lateral visual cortical functional connectivity in theta band during viewing of non-social scenes (colourful rotating toys) was positively associated with the extent of autistic symptoms later in toddlerhood. This association was strong in infants who had an elevated familial likelihood of developing autism, but non-significant in the low-likelihood group. Furthermore, among infants who later developed more autistic symptoms, this far lateral functional connectivity was seen to be stronger compared to infants who did not develop symptoms regardless of familial likelihood. We observed no association between alpha lateral connectivity, for either stimulus type, with later autism symptoms.

Consolidating our findings here with those from the previous study^7^ (which involved mostly the same infants but with data from a different experiment), we have now converging evidence in support of atypical neural organization (as reflected in both SSVEP band power discrepancy and synchronized activation between midline and peripheral lateral areas) of the visual cortex of young infants who later present with significant autistic symptoms. In the previous study^7^, the stronger lateral relative to midline SSVEP power was seen during the processing of global motion (coherent moving dots). Here, we found stronger theta but weaker gamma midline-and-peripheral lateral synchronization during non-social scene processing and stronger gamma midline-and-peripheral lateral synchronization during social scene processing.

In infants’ visual cortex, global motion laterality was found to be correlated with far-lateral gamma connectivity during processing of social scenes, but not with far-lateral theta connectivity during processing of non-social scenes. These findings pointed to the possibility that laterality of non-social theta connectivity and laterality of social gamma connectivity/global motion processing reflect independent neural processes which are both linked to autism. Finally, the main model with only a handful of predictors representing visual cortical functional connectivity, i.e., non-social theta and both social and non-social gamma, could explain almost 50% of the variability in later symptom severity of autism.

### Gamma connectivity in the visual cortex, links to global motion processing and autism

We found that the midline-to-far lateral visual cortical gamma connectivity irrespective of stimulus type was associated, albeit in different directions for social and non-social stimuli, with later autistic symptoms. These effects were only seen in the main-effect model and were not significant when moderation by the autism likelihood status was included.

Gamma rhythm activities in infancy have been linked to eye-gaze and dynamic face processing^32^, both of which have been reported to be different in autism^33-35^. Atypical facial motion processing in autism has been thought to be of perceptual nature, which could originate from altered integration of cross-feature dynamics^36^. Consistent with this, gamma rhythm has been linked to perceptual binding, needed to perceive connected parts as a single percept, and which has been shown to be present in 8-month-old infants with spatial distribution around the parieto-occipital region^37^. We found that the global motion laterality was positively linked to laterality of gamma connectivity during social scene processing, and both in turn were linked to later autistic symptoms. This may suggest that the two perceptual functions (global motion and facial motion perception) rely on some shared perceptual integrative processes that could be different in autism^38,39^. Altered unisensory and multisensory temporal integrations have been well studied in adults and children with autism^40-42^. Nevertheless, the two functions were operationalized using different analytical constructs (i.e., phase-lagged connectivity vs. SSVEP band power) each of which carries a different neurological meaning, which made the inference about the nature of the association difficult (i.e., they can be different processes relying on the same underlying mechanism or just different aspects of the same process). It is noteworthy to mention that in a previous related study^43^ we found the same global motion perception was associated with receptive language ability at 18 months (in largely the same infants), a skill that could be affected by differences in facial motion perception. Indeed, a recent study^44^ suggested the existence of a third visual cortical processing pathway specialized for social perception, on top of the well-known dorsal and ventral pathways^45^, with ascribed functionality largely consistent with our interpretation here. In particular, the midline-to-far lateral functional connectivity used in our analysis could reflect the direct connectivity between V1 and MT/V5 reported by the authors and gamma rhythm synchronization between these two areas could serve as the underlying mechanism mediating social perception through visual motion integration^44^. Our results, thus, could provide important new information about the early development of this putative third pathway as well as insights to address its autism-related questions.

We also found that gamma connectivity laterality for non-social stimuli was associated with later symptom intensity, albeit negatively. A possible explanation for both gamma connectivity results would be related to the neuronal inhibitory process, which are known to be indexed by the visual cortical gamma synchronization as seen in animal experiments^46^. The higher social gamma and lower non-social gamma connectivity associated with higher later autism symptoms might indicate an atypical balance of inhibitory processes in the visual cortex when responding to social and non-social stimuli^47^.

### Strong lateral theta connectivity in infants with later symptoms of autism

Theta oscillations (3-5 Hz) have been found predominantly within the occipital/occipitoparietal region between age 3-6 months^48,49^ and, via experimental manipulations, have been associated with infants’ emotional arousal^27,50,51^, reward experience^26^ (also in visual cortex of animal models^52^), affection-inducing cognitive processing (e.g., during exploration of novelties^22^ or a social engagement^49,53^), and sustained anticipatory attention^28,54^. While these findings stemmed from different study designs compared to ours, it is possible that the spontaneous theta connectivity arising from watching the non-social video is linked to similar processes. Specifically, the higher non-social theta connectivity in the EL-High ADOS infants might reflect increased recruitment of affect-driven attentional resources^22,54^ during the processing of the colourful rotating toy video, consistent with evidence of stronger non-social responses from other studies using a similar paradigm but with different analytical or imaging modalities (e.g., band power^53^, ERPs^55^, and fNIRS^56,57^). Furthermore, using similar social/non-social stimuli as ours, a recent study^49^ showed that in typically-developing infants of age 9 months, a reconfiguration and expansion of the cortical network synchronized in theta band^58^ occurred, making it resemble the adult’s theta network^49^. Before such reconfiguration, theta functional connectivity showed no selectivity (modulation) to either types of stimulus^49^. In line with this, the theta connectivity responses of the LL and the EL-Low-ADOS groups showed similar (flipped) normalized topographical patterns for both stimulus types. In contrast, the midline-to-far lateral theta connectivity was stronger in response to non-social relative to social stimuli in the EL-High-ADOS infants. (*Note:* No group difference was found in theta connectivity for the social stimulus.) It is possible that this finding in the EL-high-ADOS group reflects early atypical selectivity in the cortical regions (e.g., lateral occipital and posterior temporal sites^59,60^) known to be selective for social stimuli which occurred already at age 5 months^61^.

Another possible explanation for our theta result in non-social stimulus condition relates to altered cortico-subcortical functional connectivity observed in autism^62,63^, that has been found to influence sensory information processing^64-66^. Increased excitatory connectivity from the thalamus to a posterior area of temporal lobe (close to MT/V5) has been found in autistic adults^65,67^. Because the thalamus also projects to V1, an elevated connectivity to the posterior temporal lobe in autism could lead to stronger synchronization between the midline and corresponding peripheral area of the visual cortex in this group. There is evidence of thalamic regulation of cortical sensory processing (and perceptual binding) via theta-gamma^68^ and theta-beta^69^ cross-frequency coupling. Nonetheless, the specific result we saw in infants during non-social stimulus viewing needs to be explained by further investigation.

### Alpha connectivity and general attentional level to stimuli

We did not find any group or condition difference in the topographical alpha connectivity patterns. As alpha rhythm is associated with general attention and inactivity^29,70^, this may indicate that our attention coding and minimum epoch strategies worked well to ensure all infants, regardless of group, contributed data with the same level of general attentiveness to both stimulus types. Furthermore, the result also implied that the phenomena observed with both theta and gamma rhythms were not confounded by infants’ differential engagement behaviour during the session. Previous studies using similar paradigm as ours found alpha hyperconnectivity in the frontocentral scalp areas, which we did not assess here, in infants of age 14 months and found it was linked to the repetitive and restricted behaviour domain of autism^15,16^.

Finally, the link between autism and functional connectivity in general could reflect atypical cortical lateralization and asymmetry^4,71^ due to, among others, differences in early development of white matter tracts^72-75^. Yet it is notable that our results from 5-month-old infants showed the individual topographic connectivity patterns to be strongly unilateral, reflecting that the lateralization was highly individualized^76^ (not consistently to one side of the brain).

### Limitations

We did not include analysis of scalp areas associated with attention (e.g., posterior parietal, prefrontal) because the focus of our study was to follow up of our previous finding, linked to the visual cortex. Nonetheless, information about the general state of infants’ attention can be inferred from the variability of occipital alpha connectivity, which we found to be similar across experimental conditions and groups. Additionally, we have tried to minimize the risk that our analytical variables are impacted by overt attentional differences by implementing the attention coding and minimum epoch number strategies.

Our sample size suffered from high exclusion rate due to the minimum number of (≥ 90) epochs criterion which disproportionately excluded infants with high level autistic symptoms and/or an autism diagnosis. As a result, we did not have enough individuals diagnosed with autism to perform an analysis/modelling of autism diagnosis.

### Conclusion

The link between global motion and social perception has been a longstanding question in autism research^77^. Here, we illuminated one possible link between the two perceptual functions by demonstrating a correlation between the laterality of global motion processing (represented by the Global Motion Laterality Score) and laterality of gamma connectivity during social stimulus viewing (represented by the Gamma-Social Far-Connectivity). Additionally, we found another independent neural process indexed by the laterality of theta connectivity during non-social stimulus viewing which was a strong predictor of later autism. This study and our previous report^7^ converge on highlighting the atypical functional organisation within the visual cortex as an important factor in early autistic development^9^ and the potential for this brain area for future biomarker discovery.

## METHODS

### Participants

The EASE sample has been comprehensively described in past studies from our group^6,7,43^, thus we will only provide a cursory overview here. Participants were recruited via a direct phone call or adverts in paediatrics clinics around the Greater Stockholm area (for the Elevated Likelihood for Autism / EL infants) and from the Swedish Population Registry (for the Low Likelihood for Autism / LL infants). EL infants were determined from the presence of an older sibling, or any second-degree family member diagnosed with ASD. In addition to previously reported 55 infants^7^, 36^th^ month ADOS-2 scores from additional 27 infants now became available as they reached suitable age to be clinically assessed. Detailed breakdowns of analysis-level exclusion (“data attrition”) are given in **Supplementary Tables 1-2**.

### Ethics approval

All methods were performed in accordance with the relevant guidelines and regulations. Ethical permit for the EASE study was obtained in advance from the Stockholm Regional Ethics Committee.

### Informed consent

All parents or legal guardians of the participating infants signed a written informed consent on behalf of their infants for study participation at the start of study.

### Measures

#### General EEG setup

EEG recording was performed at one time-point only, when the participant was 5 months old, using the same EEG setup as reported previously^7^.

#### Visual stimuli

During the EEG session, infants watched various interchanging short video clips while sitting on their parents’ lap. A 1-minute clip showing a woman smiling and singing a Swedish nursery rhyme against a plain dark backdrop was used as our social-scene stimulus. Another 1-minute clip of two different colourful toys rotating around its axis shown one after another was used as the non-social stimulus. Both clips were played for three times in an interleaving manner, together with the form and motion stimulus videos^7^, during the 10-minute EEG session. Screenshots of both stimuli are shown in **Figs. 1a, b** and the video clips are included as **Supplementary Videos 1-2** to this manuscript.

#### Functional connectivity measure and frequency bands

Infants’ visual cortical functional connectivity was measured using dbWPLI^14^ due to its proven record for use with infants^15,16^, especially to cope with infants’ small head size that would aggravate the problems of spurious short-range connectivity. A dbWPLI score is calculated by first computing the cross-spectrum between waves measured at two different EEG electrodes and then estimating their mean phase difference (lag/lead) for a specific frequency. Mean phase difference for a range (“band”) of frequencies is calculated by averaging the scores over the frequencies. For the purposes of our study, we only take the absolute dbWPLI scores, which represent the strength of the connectivity but not the direction (i.e., leading or lagging).

Three frequency bands, adjusted for infancy^53,78^, were included in our analyses: theta (3 – 5 Hz), alpha (6 – 8 Hz), and whole gamma (20 – 60 Hz). These three brain rhythms were chosen due to their relevance to social cognitive development and autism in infants^21-25^. Therefore, together with the two stimulus types (social & non-social, see “Visual stimuli” above) we had 6 stimulus conditions (theta-social, theta-non-social, alpha-social, alpha-non-social, gamma-social, gamma-non-social) that were used in analyses.

#### Laterality score

We used the same Global Motion Laterality Score variable reported in our previous study^7^. This variable represents activation differences between midline area (presumably, corresponds to V1) and lateral area of the visual cortex^7^ and was calculated based on the T^2^-circular (T2circ) statistic^79^. The Global Motion condition corresponds to activation in response to coherent visual motion stimulus presented once every 0.5 seconds (2 Hz)^6,7^ to infants.

#### Assessment of autistic symptoms

A clinical assessment of infants’ autistic symptoms was performed at age of 36 months. As part of this assessment, ADOS-2 was administered to evaluate the severity of autistic symptoms by research-reliable clinicians. We used the total score of ADOS-2 CS as the outcome variable in a regression model (see “Analysis: Statistical modelling” below). The clinical diagnostic categorization (DSM-5) was based on a comprehensive assessment of all clinical information and consensus between two clinical experts, but as noted above only four children in the final sample received a formal DSM-5 diagnosis of autism spectrum disorder.

### Analysis

#### Coding of infants’ attention to stimuli

We performed coding of infants’ attention to the visual stimuli from the recorded infant behaviour videos taken during the sessions. For each infant, using the Datavyu^80^ software, we manually marked segments of the video where the infant either i) made gross movements (e.g., moving the whole body, an abrupt and large limb swing), or ii) looked away from the presented stimulus, where in most cases both co-occurred. The obtained time markers were then used to exclude the corresponding EEG segments from the subsequent analysis. One infant was excluded from the beginning due to having no video.

#### Exclusion of bad EEG data

Due to equipment malfunction and other causes, two infants only had less than 1 minute EEG data; the normal duration was around 10 minutes). These two infants were removed from further analysis.

#### EEG pre-processing

To pre-process the collected EEG data, we used the Harvard Automated Processing Pipeline for EEG (HAPPE)^81,82^ version 3.0 (https://github.com/PINE-Lab/HAPPE), that has been shown to provide a high data retention rate from developmental populations^83,84^. The pipeline runs on top of EEGLab^85^ and takes a raw EEG data (e.g., .RAW), processes it through various standardized steps (e.g., filtering, removing and interpolating bad EEG channels, re-referencing, segmenting, etc.) as well as applying more sophisticated techniques (e.g., 2-stage independent component analysis) to reject artefacts, and provides clean, segmented data (“epochs”) as its output^82^. Details of parameter settings used for running HAPPE in this study as well as the number of epochs produced for each processed participant’s data can be found in **Supplementary Note 1**. One participant was excluded due to having corrupted data and completely rejected by HAPPE.

#### Selection of Visual Cortex electrodes and definition of areas of interest

We inferred the EEG electrodes (within a 128-channel Geodesic Sensor Net HCGSN 130 [Electrical Geodesic Inc., OR]) that would approximately cover the entire visual cortex from past studies investigating the source localizations of EEG channels^86-88^ as well as studies of EEG functional connectivity in infants^15,16^. Twenty-four electrodes, symmetrically from left (12) and right (12) hemispheres, were selected from the occipital lobe that approximately corresponds to visual cortical areas relevant for our research questions^86,88^. The subsequent steps were generally inspired by approaches described in ref^17,18^ but adapted for use with EEG data. From this set of electrodes, we assigned two midline electrodes, Oz and POz, to define our seed area. Then, using the positional XYZ-coordinates generated by EEGLab for each electrode, we defined three areas of interest (AoIs) on each lateral side around the seed area (thus, a total of 6 AoIs) based on the Euclidean distance of each non-seed electrode to the average coordinates of the two seed electrodes. These AoIs were then assigned as areas at 1-, 2-, and 3-step distance away from the seed area, respectively, on left and right hemispheres. Please see **Fig. 1c** for a schematic diagram of these arrangements.

#### Computation of functional connectivity scores

Following the recommendations in refs^89,90^, we segmented the EEG data into 1-second 50%-overlapping epochs and, after rejecting epochs where the infant was not looking at stimuli (in more than 50% of an epoch’s length), retained only participants having at least 90 surviving epochs (equivalent to 1.5 minute of uninterrupted data). Each 1-second epoch was tapered at the edges by a Hanning window to minimize “crosstalk” between successive epochs (especially so due to the 50%-overlap) in the subsequent dbWPLI score calculation. Thereafter, we applied a Fast Fourier Transform (FFT) to obtain the power spectrum from each epoch.

Next, we proceeded with a 4-step procedure to obtain our functional connectivity variables for modelling: 1) we computed a bivariate (electrode-to-electrode) dbWPLI score, as described in ref.^14^, between each of the 22 non-seed electrodes in our “visual cortex” to each of the two seeds; 2) one single functional connectivity score for each non-seed electrode was computed by averaging the two scores from the previous step; 3) a connectivity score for an AoI was computed by averaging the scores over all non-seed electrodes belonging to that AoI; 4) these steps were repeated for each of the six experimental conditions; see “Functional connectivity measure and frequency bands” above. See **Figs. 2a** and **2b** for the plots of resulting individual and group-mean topographical dbWPLI patterns, respectively.

#### Vector-normalization and flipped topography

We applied a vector-normalization to each participant’s topographical (across the 6 AoIs) connectivity scores to focus on the relative patterns over the visual cortex and eliminate the influence of magnitude differences that can vary widely across or even within individuals, and to maintain consistency with the approach in ref^7^. This approach also had the effect of eliminating the very-high correlations (*r* ≥ .6) between the unnormalized AoI scores, which would render them practically unusable for regression modelling. Plots of these normalized topographical patterns are presented in **Figs. 2c** and **2d**.

Next, to see whether the patterns of topographical connectivity involved bilaterally strong synchronizations between V1 and the lateral extrastriate areas, i.e., if the “peak” connectivity occurred in both hemispheres or only in one, we flipped the individual topographical patterns as such that all the (normalized) maximum scores lay on the same (right, chosen arbitrarily) side of the plot. Thus, for patterns having the maximum on left hemisphere, we rotated the entire patterns around the y-axis, while those having the maximum on right hemisphere were left unchanged. This approach, which was also used in ref^7^ as a supplementary analysis, revealed a strongly unilateral nature of the topographical connectivity, with much of the group differences visible on the hemisphere where the maximum occurred (see **Figs. 2e** and **2f**). Consequently, we used these flipped patterns in all subsequent modelling and focussed only on the 3 AoIs from the (right) lateral side where the maxima are located. Although this decision removed any information about hemispheric specificity of the connectivity patterns, we considered it acceptable as it would play no role in addressing our research questions (see **Introduction** and also published as our analysis plan at https://osf.io/e8y5x).

#### Testing group differences

To infer group differences of the whole connectivity topography (all 6 AoIs included), we resorted to using permutation tests^91^ due to the highly-correlated nature of dbWPLI scores across visual cortex, even after vector-normalization and flipping. To do this, we shuffled the group labels (LL, EL-LoADOS, EL-HiADOS) and randomly assigned them among the participants, computed the difference in mean normalized dbWPLI scores for each pair of “new” groups in the 6 AoIs and 6 conditions (see “Functional connectivity measure and frequency bands”), and repeated the process for 10,000 times. The resulting distributions of normalized scores (in 6 AoIs and 6 conditions) were then tested against our data using their empirical quantiles. Consequently, this procedure performed three comparisons, one for each possible pairing of our three groups (see **Fig. 3d** and **Supplementary Fig. 2**).

To test differences in each of the 12 modelling variables (see “Statistical modelling” below) across the three participant groups, we used boxplots together with Kruskal-Wallis tests (see **Fig. 3b** and **Supplementary Fig. 1**).

#### Modelling association between functional connectivity and autism

For each frequency band (theta, alpha, gamma) and each stimulus condition (social, non-social), we took the 1-step distance AoI (on the maxima-side) scores as the “Near-Connectivity” variable and the 3-step distance AoI scores (on the maxima-side) as the “Far-Connectivity” variable, giving us a total of 12 modelling variables. We skipped using the 2-step distance AoI scores to further minimize any possible “crosstalk” between neighbouring scalp regions. For simplicity, we do not report the ratio of far-to-near connectivity, which we originally planned to do (https://osf.io/e8y5x), because eventually it did not provide any additional information beyond what can be provided by the Far- and Near-Connectivity combined. The resulting correlations among these 12 variables were more amenable (see **Supplementary Note 1**) for fitting simple ordinary least square (OLS) regression models (see below).

Subsequently, a non-parametric variable selection procedure based on cross-validation^55,92,93^ was run to identify a subset of the 12 variables that best explains the variability in ADOS-2 total score. A collection of all 4,096 (= 2^12^) possible subsets of the 12 variables, including the empty set and full set, was generated and then an OLS linear regression model with ADOS-2 total score as the outcome was fitted on each of these subsets as regressors. For each fitted model, we performed a 10-fold cross-validation (CV) with 10 repetitions (each time generating a different random 9:1 split of the dataset) to obtain an average coefficient of determination (*R*^2^) score, selected 100 models with the best *R*^2^ scores, and ranked the frequency of occurrences of each variable in these 100 models. The top 5 variables were then fitted in the final linear regression model for inferring the association with ADOS-2 total score.

#### Relation between functional connectivity and global motion processing

To infer whether variance (of ADOS-2 total score) explained by the dbWPLI variables can be explained by the Global Motion Laterality Score, we added the latter to the existing linear model of ADOS-2 total score vs. functional connectivity (see “Modelling association between functional connectivity and autism”) as an extra predictor. Then, we (i) noted how the sum-of-square (SS) of each dbWPLI predictor changed and (ii) tested whether the resulting increase (if any) of the model’s *R*^2^ is significant. For (i), we performed an analysis of variance (ANOVA) on the model before and after addition of the extra predictor, and for (ii) a nested likelihood ratio test on the same two models was done. A substantial overlap in variances explained by Global Motion Laterality Score and any of the dbWPLI predictors would point to a possibility that the lateral activation we saw in the previous study of global motion processing^7^ has a basis in and/or an implication for functional connectivity in the brain rhythms studied here. Finally, to better understand the nature of this hypothesized relation between global motion processing and visual cortical functional connectivity in the three brain rhythms, we fitted an OLS linear regression of Global Motion Laterality Score regressing on dbWPLI predictors. Before the model-fitting, we again ran the same non-parametric variable selection procedure as before (see “Modelling association between functional connectivity and autism”), but now with Global Motion Laterality Score as the outcome, to trim down the number of plausible dbWPLI predictors to include in the model from 12 to only 5. The eventual model fitting included only the selected 5 dbWPLI predictors and no covariates, because sex, age (at 5 months), and likelihood for autism status had been regressed out from all dbWPLI variables and Global Motion Laterality score.

## Data availability

The data that support the findings of this study are not openly available due to reasons of sensitivity and are available from the corresponding author upon reasonable request.

## Code availability

All code is available upon request.

## Acknowledgments

This study was funded by Riksbankens Jubileumsfond, the Knut and Alice Wallenberg Foundations and the Innovative Medicines Initiative 2 Joint Undertaking under grant agreement No 777394. This Joint Undertaking receives support from the European Union’s Horizon 2020 research and innovation program and EFPIA and AUTISM SPEAKS, Autistica, SFARI. The work leading to these results was also supported by funds from the European Commission (H2020 project CANDY; Grant No. 847818); and from Marie S. Curie Actions SAPIENS Project (Grant MSCA-ITN-2018 No. 814302). The funders had no role in the design of the study; in the collection, analyses, or interpretation of data; in the writing of the manuscript, or in the decision to publish the results.

## Competing interests

The authors declare no competing interests. S.B. discloses that he has in the last 5 years acted as an author, consultant, or lecturer for Medice and Roche. He receives royalties for textbooks and diagnostic tools from Hogrefe (ADOS-2, ADI-R, SRS-2, SCQ), Kohlham-mer, and UTB.

## Notes

### Competing Interest Statement

The authors have declared no competing interest.

## References

1 Courchesne, E. et al. Unusual brain growth patterns in early life in patients with autistic disorder: an MRI study. Neurology 57, 245–254 (2001).

2 Shen, M. D. et al. Early brain enlargement and elevated extra-axial fluid in infants who develop autism spectrum disorder. Brain 136, 2825–2835 (2013).

3 Emerson, R. W. et al. Functional neuroimaging of high-risk 6-month-old infants predicts a diagnosis of autism at 24 months of age. Science translational medicine 9, eaag2882 (2017).

4 Postema, M. C. et al. Altered structural brain asymmetry in autism spectrum disorder in a study of 54 datasets. Nature communications 10, 1–12 (2019).

5 Hazlett, H. C. et al. Early brain development in infants at high risk for autism spectrum disorder. Nature 542, 348–351 (2017).

6 Nyström, P., Jones, E., Darki, F., Bölte, S. & Falck-Ytter, T. Atypical topographical organization of global form and motion processing in 5-month-old infants at risk for Autism. Journal of autism and developmental disorders 51, 364–370 (2021).

7 Hardiansyah, I. et al. Global motion processing in infants’ visual cortex and the emergence of autism. Communications Biology 6, 1–10 (2023).

8 Gandal, M. J. et al. Broad transcriptomic dysregulation occurs across the cerebral cortex in ASD. Nature, 1–8 (2022).

9 Falck-Ytter, T. & Bussu, G. The sensory-first account of autism. Neuroscience and Biobehavioral Reviews 153 (2023).

10 Belmonte, M. K. et al. Autism and abnormal development of brain connectivity. Journal of Neuroscience 24, 9228–9231 (2004).

11 Uddin, L. Q., Supekar, K. & Menon, V. Reconceptualizing functional brain connectivity in autism from a developmental perspective. Frontiers in human neuroscience 7, 458 (2013).

12 O’Reilly, C., Lewis, J. D. & Elsabbagh, M. Is functional brain connectivity atypical in autism? A systematic review of EEG and MEG studies. PloS one 12, e0175870 (2017).

13 Müller, R.-A. & Fishman, I. Brain connectivity and neuroimaging of social networks in autism. Trends in cognitive sciences 22, 1103–1116 (2018).

14 Vinck, M., Oostenveld, R., Van Wingerden, M., Battaglia, F. & Pennartz, C. M. An improved index of phase-synchronization for electrophysiological data in the presence of volume-conduction, noise and sample-size bias. Neuroimage 55, 1548–1565 (2011).

15 Haartsen, R., Jones, E. J., Orekhova, E. V., Charman, T. & Johnson, M. H. Functional EEG connectivity in infants associates with later restricted and repetitive behaviours in autism; a replication study. Translational Psychiatry 9, 66 (2019).

16 Orekhova, E. V. et al. EEG hyper-connectivity in high-risk infants is associated with later autism. Journal of neurodevelopmental disorders 6, 1–11 (2014).

17 Sepulcre, J., Sabuncu, M. R., Yeo, T. B., Liu, H. & Johnson, K. A. Stepwise connectivity of the modal cortex reveals the multimodal organization of the human brain. Journal of Neuroscience 32, 10649–10661 (2012).

18 den Bakker, H., et al. Abnormal coherence and sleep composition in children with Angelman syndrome: a retrospective EEG study. Molecular autism 9, 1–12 (2018).

19 Peters, J. M. et al. Brain functional networks in syndromic and non-syndromic autism: a graph theoretical study of EEG connectivity. BMC medicine 11, 1–16 (2013).

20 Jones, E. J. et al. Infant EEG theta modulation predicts childhood intelligence. Scientific reports 10, 11232 (2020).

21 Mittag, M., Larson, E., Taulu, S., Clarke, M. & Kuhl, P. K. Reduced theta sampling in infants at risk for dyslexia across the sensitive period of native phoneme learning. International Journal of Environmental Research and Public Health 19, 1180 (2022).

22 Orekhova, E., Stroganova, T., Posikera, I. & Elam, M. EEG theta rhythm in infants and preschool children. Clinical neurophysiology 117, 1047–1062 (2006).

23 Larrain-Valenzuela, J. et al. Theta and alpha oscillation impairments in autistic spectrum disorder reflect working memory deficit. Scientific reports 7, 14328 (2017).

24 Orekhova, E. V. et al. Gamma oscillations point to the role of primary visual cortex in atypical motion processing in autism. Plos one 18, e0281531 (2023).

25 Sperdin, H. F. et al. Early alterations of social brain networks in young children with autism. Elife 7, e31670 (2018).

26 Maulsby, R. L. An illustration of emotionally evoked theta rhythm in infancy: Hedonic hypersynchrony. Electroencephalography and Clinical Neurophysiology 31, 157–165 (1971).

27 Posikera, I., Stroganova, T., Zhurba, L. & Timonina, O. Electroencephalographic correlates of positive emotional reactions in children in the first year of life. Zhurnal Nevropatologii i Psikhiatrii Imeni SS Korsakova (Moscow, Russia: 1952) 86, 1485–1491 (1986).

28 Orekhova, E. V., Stroganova, T. A. & Posikera, I. N. Theta synchronization during sustained anticipatory attention in infants over the second half of the first year of life. International Journal of Psychophysiology 32, 151–172 (1999).

29 Stroganova, T. A., Orekhova, E. V. & Posikera, I. N. EEG alpha rhythm in infants. Clinical neurophysiology 110, 997–1012 (1999).

30 Gotham, K., Pickles, A. & Lord, C. Standardizing ADOS scores for a measure of severity in autism spectrum disorders. Journal of autism and developmental disorders 39, 693–705 (2009).

31 Lord, C., et al. Autism diagnostic observation schedule–2nd edition (ADOS-2). Los Angeles, CA: Western Psychological Corporation 284 (2012).

32 Grossmann, T., Johnson, M. H., Farroni, T. & Csibra, G. Social perception in the infant brain: gamma oscillatory activity in response to eye gaze. Social cognitive and affective neuroscience 2, 284–291 (2007).

33 Nomi, J. S. & Uddin, L. Q. Face processing in autism spectrum disorders: From brain regions to brain networks. Neuropsychologia 71, 201–216 (2015).

34 Reisinger, D. L. et al. Atypical social attention and emotional face processing in autism spectrum disorder: Insights from face scanning and pupillometry. Frontiers in Integrative Neuroscience 13, 76 (2020).

35 Stuart, N., Whitehouse, A., Palermo, R., Bothe, E. & Badcock, N. Eye gaze in autism spectrum disorder: a review of neural evidence for the eye avoidance hypothesis. Journal of Autism and Developmental Disorders 53, 1884–1905 (2023).

36 Shah, P., Bird, G. & Cook, R. Face processing in autism: Reduced integration of cross-feature dynamics. cortex 75, 113–119 (2016).

37 Csibra, G., Davis, G., Spratling, M. & Johnson, M. Gamma oscillations and object processing in the infant brain. Science 290, 1582–1585 (2000).

38 Thye, M. D., Bednarz, H. M., Herringshaw, A. J., Sartin, E. B. & Kana, R. K. The impact of atypical sensory processing on social impairments in autism spectrum disorder. Developmental cognitive neuroscience 29, 151–167 (2018).

39 O’Brien, J., Spencer, J., Girges, C., Johnston, A. & Hill, H. Impaired perception of facial motion in autism spectrum disorder. PLoS One 9, e102173 (2014).

40 Nakano, T., Ota, H., Kato, N. & Kitazawa, S. Deficit in visual temporal integration in autism spectrum disorders. Proceedings of the Royal Society B: Biological Sciences 277, 1027–1030 (2010).

41 Stevenson, R. A. et al. Multisensory temporal integration in autism spectrum disorders. Journal of Neuroscience 34, 691–697 (2014).

42 Meilleur, A., Foster, N. E., Coll, S.-M., Brambati, S. M. & Hyde, K. L. Unisensory and multisensory temporal processing in autism and dyslexia: A systematic review and meta-analysis. Neuroscience & Biobehavioral Reviews 116, 44–63 (2020).

43 Hedenius, M., Hardiansyah, I. & Falck-Ytter, T. Visual global processing and subsequent verbal and non-verbal development: An EEG study of infants at elevated versus low likelihood for autism spectrum disorder. Journal of autism and developmental disorders, 1–10 (2022).

44 Pitcher, D. & Ungerleider, L. G. Evidence for a third visual pathway specialized for social perception. Trends in Cognitive Sciences 25, 100–110 (2021).

45 Mishkin, M., Ungerleider, L. G. & Macko, K. A. Object vision and spatial vision: two cortical pathways. Trends in neurosciences 6, 414–417 (1983).

46 Veit, J., Hakim, R., Jadi, M. P., Sejnowski, T. J. & Adesnik, H. Cortical gamma band synchronization through somatostatin interneurons. Nature neuroscience 20, 951–959 (2017).

47 Klin, A., Shultz, S. & Jones, W. Social visual engagement in infants and toddlers with autism: early developmental transitions and a model of pathogenesis. Neuroscience & Biobehavioral Reviews 50, 189–203 (2015).

48 Marshall, P. J., Bar-Haim, Y. & Fox, N. A. Development of the EEG from 5 months to 4 years of age. Clinical neurophysiology 113, 1199–1208 (2002).

49 van der Velde, B., White, T. & Kemner, C. The emergence of a theta social brain network during infancy. Neuroimage 240, 118298 (2021).

50 Futagi, Y., Ishihara, T., Tsuda, K., Suzuki, Y. & Goto, M. Theta rhythms associated with sucking, crying, gazing and handling in infants. Electroencephalography and clinical neurophysiology 106, 392–399 (1998).

51 Lehtonen, J., Könönen, M., Purhonen, M., Partanen, J. & Saarikoski, S. The effects of feeding on the electroencephalogram in 3-and 6-month-old infants. Psychophysiology 39, 73–79 (2002).

52 Zold, C. L. & Shuler, M. G. H. Theta oscillations in visual cortex emerge with experience to convey expected reward time and experienced reward rate. Journal of Neuroscience 35, 9603–9614 (2015).

53 Jones, E. J., Venema, K., Lowy, R., Earl, R. K. & Webb, S. J. Developmental changes in infant brain activity during naturalistic social experiences. Developmental psychobiology 57, 842–853 (2015).

54 Stroganova, T. A., Orekhova, E. V. & Posikera, I. N. Externally and internally controlled attention in infants: an EEG study. International Journal of Psychophysiology 30, 339–351 (1998).

55 Gui, A. et al. Attentive brain states in infants with and without later autism. Translational psychiatry 11, 196 (2021).

56 Lloyd-Fox, S. et al. Reduced neural sensitivity to social stimuli in infants at risk for autism. Proceedings of the Royal Society B: Biological Sciences 280, 20123026 (2013).

57 Lloyd - Fox, S., et al. Cortical responses before 6 months of life associate with later autism. European Journal of Neuroscience 47, 736–749 (2018).

58 Johnson, E. L. et al. A rapid theta network mechanism for flexible information encoding. Nature Communications 14, 2872 (2023).

59 Johnson, M. H. et al. The emergence of the social brain network: Evidence from typical and atypical development. Development and psychopathology 17, 599–619 (2005).

60 Haan, M. d., Pascalis, O. & Johnson, M. H. Specialization of neural mechanisms underlying face recognition in human infants. Journal of cognitive neuroscience 14, 199–209 (2002).

61 Grossmann, T. & Johnson, M. H. The development of the social brain in human infancy. European Journal of Neuroscience 25, 909–919 (2007).

62 Park, B.-y., et al. Differences in subcortico-cortical interactions identified from connectome and microcircuit models in autism. Nature communications 12, 2225 (2021).

63 Mizuno, A., Villalobos, M. E., Davies, M. M., Dahl, B. C. & Müller, R.-A. Partially enhanced thalamocortical functional connectivity in autism. Brain research 1104, 160–174 (2006).

64 Colich, N. L. et al. Atypical neural processing of ironic and sincere remarks in children and adolescents with autism spectrum disorders. Metaphor and symbol 27, 70–92 (2012).

65 Woodward, N. D., Giraldo-Chica, M., Rogers, B. & Cascio, C. J. Thalamocortical dysconnectivity in autism spectrum disorder: an analysis of the autism brain imaging data exchange. Biological Psychiatry: Cognitive Neuroscience and Neuroimaging 2, 76–84 (2017).

66 McFadyen, J., Dolan, R. J. & Garrido, M. I. The influence of subcortical shortcuts on disordered sensory and cognitive processing. Nature Reviews Neuroscience 21, 264–276 (2020).

67 Nair, A., Treiber, J. M., Shukla, D. K., Shih, P. & Müller, R.-A. Impaired thalamocortical connectivity in autism spectrum disorder: a study of functional and anatomical connectivity. Brain 136, 1942–1955 (2013).

68 Ribary, U., Doesburg, S. & Ward, L. Unified principles of thalamo-cortical processing: the neural switch. Biomedical Engineering Letters 7, 229–235 (2017).

69 Malekmohammadi, M., Elias, W. J. & Pouratian, N. Human thalamus regulates cortical activity via spatially specific and structurally constrained phase-amplitude coupling. Cerebral Cortex 25, 1618–1628 (2015).

70 Xie, W., Mallin, B. M. & Richards, J. E. Development of infant sustained attention and its relation to EEG oscillations: an EEG and cortical source analysis study. Developmental science 21, e12562 (2018).

71 Kong, X. Z. et al. Mapping brain asymmetry in health and disease through the ENIGMA consortium. Human brain mapping 43, 167–181 (2022).

72 Wolff, J. J. et al. Differences in white matter fiber tract development present from 6 to 24 months in infants with autism. American journal of Psychiatry 169, 589–600 (2012).

73 Wolff, J. J. et al. Neural circuitry at age 6 months associated with later repetitive behavior and sensory responsiveness in autism. Molecular autism 8, 1–12 (2017).

74 Piven, J., Elison, J. T. & Zylka, M. J. Toward a conceptual framework for early brain and behavior development in autism. Molecular Psychiatry 22, 1385–1394 (2017).

75 Song, J. W. et al. Asymmetry of white matter pathways in developing human brains. Cerebral cortex 25, 2883–2893 (2015).

76 Floris, D. L. et al. Atypical brain asymmetry in autism—a candidate for clinically meaningful stratification. Biological Psychiatry: Cognitive Neuroscience and Neuroimaging 6, 802–812 (2021).

77 Van der Hallen, R., Manning, C., Evers, K. & Wagemans, J. Global motion perception in autism spectrum disorder: a meta-analysis. Journal of autism and developmental disorders 49, 4901–4918 (2019).

78 Saby, J. N. & Marshall, P. J. The utility of EEG band power analysis in the study of infancy and early childhood. Developmental neuropsychology 37, 253–273 (2012).

79 Victor, J. D. & Mast, J. A new statistic for steady-state evoked potentials. Electroencephalography and clinical neurophysiology 78, 378–388 (1991).

80 Team, D. Datavyu: A video coding tool. Databrary Project, New York University. URL http://datavyu.org (2014).

81 Gabard-Durnam, L. J., Mendez Leal, A. S., Wilkinson, C. L. & Levin, A. R. The Harvard Automated Processing Pipeline for Electroencephalography (HAPPE): standardized processing software for developmental and high-artifact data. Frontiers in neuroscience 12, 97 (2018).

82 Monachino, A., Lopez, K., Pierce, L. & Gabard-Durnam, L. The HAPPE plus event-related (HAPPE+ ER) software: a standardized processing pipeline for event-related potential analyses. BioRxiv, 2021.2007. 2002.450946 (2021).

83 Auger, E., Berry-Kravis, E. M. & Ethridge, L. E. Independent evaluation of the harvard automated processing pipeline for Electroencephalography 1.0 using multi-site EEG data from children with Fragile X Syndrome. Journal of Neuroscience Methods 371, 109501 (2022).

84 Peck, F. C. et al. Prediction of autism spectrum disorder diagnosis using nonlinear measures of language-related EEG at 6 and 12 months. Journal of neurodevelopmental disorders 13, 1–13 (2021).

85 Brunner, C., Delorme, A. & Makeig, S. Eeglab–an open source matlab toolbox for electrophysiological research. Biomedical Engineering/Biomedizinische Technik 58, 000010151520134182 (2013).

86 Scrivener, C. L. & Reader, A. T. Variability of EEG electrode positions and their underlying brain regions: visualizing gel artifacts from a simultaneous EEG - fMRI dataset. Brain and Behavior 12, e2476 (2022).

87 Shirazi, S. Y. & Huang, H. J. More reliable EEG electrode digitizing methods can reduce source estimation uncertainty, but current methods already accurately identify brodmann areas. Frontiers in neuroscience 13, 1159 (2019).

88 Koessler, L. et al. Automated cortical projection of EEG sensors: anatomical correlation via the international 10–10 system. Neuroimage 46, 64–72 (2009).

89 Haartsen, R., van der Velde, B., Jones, E. J., Johnson, M. H. & Kemner, C. Using multiple short epochs optimises the stability of infant EEG connectivity parameters. Scientific Reports 10, 12703 (2020).

90 Haartsen, R., Charman, T., Pasco, G., Johnson, M. H. & Jones, E. J. Modulation of EEG theta by naturalistic social content is not altered in infants with family history of autism. Scientific Reports 12, 20758 (2022).

91 Collingridge, D. S. A primer on quantitized data analysis and permutation testing. Journal of mixed methods research 7, 81–97 (2013).

92 Tye, C. et al. Understanding the nature of face processing in early autism: a prospective study. Journal of Psychopathology and Clinical Science 131, 542 (2022).

93 Snaedal, J. et al. Diagnostic accuracy of statistical pattern recognition of electroencephalogram registration in evaluation of cognitive impairment and dementia. Dementia and geriatric cognitive disorders 34, 51–60 (2012).

