## Supplementary Information for "Functional connectivity in infants’ visual cortex: Links to motion processing and autism"

**Supplementary Table 1:** Analysis-level exclusions for functional connectivity calculation

**Supplementary Table 2:** Analysis-level exclusions for global motion laterality analysis

**Supplementary Table 3:** Descriptives of infants in functional connectivity dataset

**Supplementary Table 4:** Descriptives of infants in global motion laterality dataset

**Supplementary Table 5:** Results of statistical tests and models presented in the main text and Supplementary Information

**Supplementary Figure 1:** Group analysis of dbWPLI Near & Far Connectivity variables in six experimental conditions

**Supplementary Figure 2:** Permutation tests of across-group differences in five experimental conditions based on the flipped topographies

**Supplementary Figure 3:** Across-stimulus topographic connectivity differences in each frequency band and group

**Supplementary Table 1 | Analysis-level exclusions for functional connectivity calculation**

| | **Sample size** |
|---|---|
| Initial sample | 119† |
| No video coding, no EEG data | -2 |
| Sample having EEG & video coding | 117 |
| Bad EEG data & rejected by HAPPE for preprocessing | -3 |
| Sample pre-processed by HAPPE | 114 |
| Exclusion due to no looking to stimuli | -12 |
| Sample going into analysis stage | 102 |
| Exclusion due to the 90-epoch threshold | -34 |
| **Final sample for computing dbWPLI variables** | **69** |
| Sub-sample with ADOS-2 CSS at 36 mo. | 59 |
| Sub-sample with Global Motion Laterality Score & ADOS-2 CSS at 36 mo.‡ | 43 |

†The same participants as in ref.[1] plus 27 new infants, after excluding ADHD-only cases

‡Crf. Supplementary Table 2

**Supplementary Table 2 | Analysis-level exclusions for global motion laterality analysis[1]**

| | **Sample size** |
|---|---|
| Initial sample | 119† |
| No EEG data | -1 |
| Sample having EEG | 118 |
| Bad EEG data | -23 |
| **Final sample for computing Global Motion Laterality Score** | **95** |
| Sub-sample with ADOS-2 CSS at 36 mo. | 82 |
| Sub-sample with Global Motion Laterality Score & ADOS-2 CSS at 36 mo.‡ | 43 |

†The same participants as in ref.[1] plus 22 additional infants

‡Crf. Supplementary Table 1

**Supplementary Table 3 | Descriptives of infants in functional connectivity dataset**

| ***EEG recording at 5 months*** | **EL** | | **LL** |
|---|---|---|---|
| N | | 49 | 20 |
| Female | | 31 | 9 |
| Age in months (mean ± s.d.) | | 5.51 ± 0.66 | 5.33 ± 0.44 |
| MSEL* Early Learning Composite Score | | 96.9 ± 11.1 | 97.9 ± 10.0 |
| ***ADOS-2 at 36 months*** | **EL High-ADOS** | **EL Low-ADOS** | **LL** |
| N | 11 | 28 | 20 |
| Female | 4 | 17 | 11 |
| Age in months (mean ± s.d.) | 37.80 ± 1.95 | 37.85 ± 2.75 | 37.65 ± 3.02 |

*Mullen Scales of Early Learning (MSEL)[2]

**Supplementary Table 4 | Descriptives of infants in global motion laterality dataset[1]**

| ***EEG recording at 5 months*** | **EL** | | **LL** |
|---|---|---|---|
| N | | 71 | 24 |
| Female | | 35 | 15 |
| Age in months (mean ± s.d.) | | 5.44 ± 0.72 | 5.39 ± 0.62 |
| MSEL* Early Learning Composite Score | | 96.3 ± 10.3 | 102.1 ± 7.4 |
| ***ADOS-2 at 36 months*** | **EL High-ADOS** | **EL Low-ADOS** | **LL** |
| N | 19 | 44 | 19 |
| Female | 6 | 26 | 12 |
| Age in months (mean ± s.d.) | 38.10 ± 2.65 | 37.98 ± 2.84 | 37.46 ± 2.84 |

*Mullen Scales of Early Learning (MSEL)[2]

**Supplementary Table 5 | Results of statistical tests and models presented in the main text, Extended Data, and in Supplementary Information**

All analyses presented in the manuscript were performed using standard functions/scripts in R language. All *p*-values are <u>two-tailed</u>.

***Abbreviations:*** FC_ = Far-Connectivity; NC_ = Near-Connectivity; [Band]NS = NonSocial stimuli; [Band]S = Social stimuli; dbWPLI = debiased Weighted Phase Lag Index; GMLS = Global Motion Laterality Score; EL = Elevated Likelihood for ASD; LL = Low Likelihood for ASD; LR = Likelihood Ratio; npar = no. of model parameters; LogLik = Log-Likelihood; df = degrees of freedom; GFLS = Global Form Laterality Score; LMLS = Local Motion Laterality Score; LFLS = Local Form Laterality Score

| Category | Section in Main Text | Referred to in Display Item or on Manus. Page | Test / Model | Parameter estimates | *N* | *p* |
|---|---|---|---|---|---|---|
| Main result | **Results:** *Association between infants' visual cortical functional connectivity and autistic symptoms in toddlerhood* | Fig. 3a | Bivariate correlation (Pearson's *r*) | All: *r*(FC_ThetaNS x ADOS2) = .50<br>EL: *r*(FC_ThetaNS x ADOS2) = .64<br>LL: *r*(FC_ThetaNS x ADOS2) = -.38 | 59<br>39<br>20 | < .001<br>< .001<br>.095 |
| Main result | **Results:** *Association between infants' visual cortical functional connectivity and autistic symptoms in toddlerhood* | Fig. 3a | Main-effect model of ADOS2 ~ dbWPLI (OLS regression) | <u>**Regression coefficient [Effect size]**</u><br>*β*(sex_Male) = .87 [part-□$^2$ = .077]<br>*β*(age) = -.02 [part-□$^2$ = .050]<br>*β*(status_LL) = -.98 [part-□$^2$ = .087]<br>*β*(F_ThetaNS) = 1.12 [part-□$^2$ = .229]<br>*β*(F_GammaNS) = -.59 [part-□$^2$ = .076]<br>*β*(F_GammaS) = .60 [part-□$^2$ = .085]<br>*β*(N_ThetaNS) = .32 [part-□$^2$ = .028]<br>*β*(N_AlphaS) = .36 [part-□$^2$ = .047]<br>$R^2$ = .440 | 59 | <br>.047<br>.111<br>.034<br>< .001<br>.047<br>.036<br>.234<br>.124 |
| Main result | **Results:** *Association between infants' visual cortical functional connectivity and autistic symptoms in toddlerhood* | Fig. 3a | Interaction-effect model of ADOS2 ~ dbWPLI*status (OLS regression) | <u>**Regression coefficient [Effect size]**</u><br>*β*(sex_Male) = .71 [part-□$^2$ = .054]<br>*β*(age) = -.02 [part-□$^2$ = .036]<br>*β*(status_LL) = -.98 [part-□$^2$ = .085]<br>*β*(F_ThetaNS) = 1.18 [part-□$^2$ = .233]<br>*β*(F_GammaNS) = -.52 [part-□$^2$ = .058]<br>*β*(F_GammaS) = .45 [part-□$^2$ = .043]<br>*β*(F_ThetaNS*status_LL) = -1.91 [part-□$^2$ = .135]<br>*β*(F_GammaNS*status_LL) = .41 [part-□$^2$ = .005]<br>*β*(F_GammaS*status_LL) = -.16 [part-□$^2$ = .001]<br>$R^2$ = .493 | 59 | <br>.100<br>.181<br>.029<br>< .001<br>.079<br>.158<br>.008<br>.634<br>.823 |
| Main result | **Results:** *Association between infants' visual cortical functional connectivity and autistic* | Fig. 3b,<br>Ext. Data Fig. 1 | Across-group difference test of dbWPLI variables (Wilcoxon test + FDR correction)<br>*<u>Notes:</u>*<br>[i]EL-High ADOS | <u>**FC_ThetaNS, effect size (Cohen's *d*)**</u><br>EL Hi-ADOS – EL Lo-ADOS: *d* = 1.61<br>EL Hi-ADOS – LL: *d* = 1.57<br>EL Lo-ADOS – LL: *d* = -.14<br><br><u>**FC_ThetaS, effect size (Cohen's *d*)**</u><br>EL Hi-ADOS – EL Lo-ADOS: *d* = -.09<br>EL Hi-ADOS – LL: *d* = .44 | 59<br>11[i)]<br>29[ii)]<br>20[iii)] | <br>004<br>.008<br>.750<br><br><br>.939<br>.522 |

| | | | | | | |
|---|---|---|---|---|---|---|
| | *symptoms in toddlerhood* | | [ii]EL-Low ADOS<br>[iii]LL | EL Lo-ADOS – LL: $d$ = .46<br><br>**FC_GammaNS, effect size (Cohen's *d*)**<br>EL Hi-ADOS – EL Lo-ADOS: $d$ = .42<br>EL Hi-ADOS – LL: $d$ = .55<br>EL Lo-ADOS – LL: $d$ = .12<br><br>**FC_GammaS, effect size (Cohen's *d*)**<br>EL Hi-ADOS – EL Lo-ADOS: $d$ = .73<br>EL Hi-ADOS – LL: $d$ = .66<br>EL Lo-ADOS – LL: $d$ = .02<br><br>**FC_AlphaNS, effect size (Cohen's *d*)**<br>EL Hi-ADOS – EL Lo-ADOS: $d$ = .43<br>EL Hi-ADOS – LL: $d$ = 1.06<br>EL Lo-ADOS – LL: $d$ = .53<br><br>**FC_AlphaS, effect size (Cohen's *d*)**<br>EL Hi-ADOS – EL Lo-ADOS: $d$ = .19<br>EL Hi-ADOS – LL: $d$ = .20<br>EL Lo-ADOS – LL: $d$ = .03<br><br>**NC_ThetaNS, effect size (Cohen's *d*)**<br>EL Hi-ADOS – EL Lo-ADOS: $d$ = -.98<br>EL Hi-ADOS – LL: $d$ = -.91<br>EL Lo-ADOS – LL: $d$ = -.06<br><br>**NC_ThetaS, effect size (Cohen's *d*)**<br>EL Hi-ADOS – EL Lo-ADOS: $d$ = -.15<br>EL Hi-ADOS – LL: $d$ = -.17<br>EL Lo-ADOS – LL: $d$ = .02<br><br>**NC_GammaNS, effect size (Cohen's *d*)**<br>EL Hi-ADOS – EL Lo-ADOS: $d$ = -.34<br>EL Hi-ADOS – LL: $d$ = -.01<br>EL Lo-ADOS – LL: $d$ = .36<br><br>**NC_GammaS, effect size (Cohen's *d*)**<br>EL Hi-ADOS – EL Lo-ADOS: $d$ = -.65<br>EL Hi-ADOS – LL: $d$ = -.66<br>EL Lo-ADOS – LL: $d$ = .07<br><br>**NC_AlphaNS, effect size (Cohen's *d*)**<br>EL Hi-ADOS – EL Lo-ADOS: $d$ = -.52<br>EL Hi-ADOS – LL: $d$ = -.37<br>EL Lo-ADOS – LL: $d$ = .25<br><br>**NC_AlphaS, effect size (Cohen's *d*)**<br>EL Hi-ADOS – EL Lo-ADOS: $d$ = .03<br>EL Hi-ADOS – LL: $d$ = -.12<br>EL Lo-ADOS – LL: $d$ = -.16 | | .446<br><br><br>.545<br>.615<br>.860<br><br><br>.508<br>.508<br>.860<br><br><br>.761<br>.527<br>.761<br><br><br>.893<br>.761<br>.893<br><br><br>.022<br>.076<br>.835<br><br><br>.918<br>.833<br>.917<br><br><br>.860<br>.833<br>.508<br><br><br>.508<br>.508<br>.860<br><br><br>.761<br>.893<br>.893<br><br><br>.893<br>.893<br>.893 |
| Main result | **Results:**<br>*Association between infants' visual cortical functional connectivity and autistic symptoms in toddlerhood* | Figs. 3c-d<br>Ext. Data Fig. 2 | Across-group difference test of flipped, norm. AoI connectivity (Permutation test + FDR correct.)<br>*Notes:*<br>[i]EL-High ADOS<br>[ii]EL-Low ADOS<br>[iii]LL | **ThetaNS, mean difference [effect size]**<br>* *EL Hi-ADOS – EL Lo-ADOS*<br>Step-3L: $M$ = -.020 [$d$ = -.81]<br>Step-2L: $M$ = -.056 [$d$ = -2.36]<br>Step-1L: $M$ = -.018 [$d$ = -.74]<br>Step-1R: $M$ = -.094 [$d$ = -2.41]<br>Step-2R: $M$ = -.046 [$d$ = -1.52]<br>Step-3R: $M$ = .234 [$d$ = 4.16]<br>* *EL Hi-ADOS – LL*<br>Step-3L: $M$ = -.014 [$d$ = -.55] | 59<br>11[i]<br>28[ii]<br>20[iii] | <br><br>.445<br>.050<br>.449<br>.050<br>.264<br>< .001<br><br>.490 |

| | | | | | | |
|---|---|---|---|---|---|---|
| | | | | Step-2L: *M* = -.060 [*d* = -2.44] | | .050 |
| | | | | Step-1L: *M* = -.018 [*d* = -.68] | | .449 |
| | | | | Step-1R: *M* = -.100 [*d* = -2.44] | | .050 |
| | | | | Step-2R: *M* = -.025 [*d* = -.78] | | .446 |
| | | | | Step-3R: *M* = .217 [*d* = 3.67] | | .004 |
| | | | | * *EL Lo-ADOS – LL* | | |
| | | | | Step-3L: *M* = .006 [*d* = .29] | | .492 |
| | | | | Step-2L: *M* = -.004 [*d* = -.22] | | .492 |
| | | | | Step-1L: *M* = .000 [*d* = .02] | | .498 |
| | | | | Step-1R: *M* = -.007 [*d* = -.22] | | .492 |
| | | | | Step-2R: *M* = .021 [*d* = .86] | | .446 |
| | | | | Step-3R: *M* = -.017 [*d* = -.36] | | .492 |
| | | | | **ThetaS, mean difference [effect size]** | | |
| | | | | * *EL Hi-ADOS – EL Lo-ADOS* | | |
| | | | | Step-3L: *M* = .004 [*d* = .17] | | .491 |
| | | | | Step-2L: *M* = -.002 [*d* = -.09] | | .498 |
| | | | | Step-1L: *M* = -.002 [*d* = -.08] | | .498 |
| | | | | Step-1R: *M* = -.015 [*d* = -.45] | | .492 |
| | | | | Step-2R: *M* = .026 [*d* = .93] | | .446 |
| | | | | Step-3R: *M* = -.012 [*d* = -.26] | | .492 |
| | | | | * *EL Hi-ADOS – LL* | | |
| | | | | Step-3L: *M* = -.029 [*d* = -1.09] | | .446 |
| | | | | Step-2L: *M* = -.021 [*d* = -.81] | | .446 |
| | | | | Step-1L: *M* = .011 [*d* = .43] | | .492 |
| | | | | Step-1R: *M* = -.014 [*d* = -.38] | | .492 |
| | | | | Step-2R: *M* = .005 [*d* = .16] | | .492 |
| | | | | Step-3R: *M* = .048 [*d* = 1.01] | | .446 |
| | | | | * *EL Lo-ADOS – LL* | | |
| | | | | Step-3L: *M* = -.033 [*d* = -1.60] | | .245 |
| | | | | Step-2L: *M* = -.018 [*d* = -.91] | | .446 |
| | | | | Step-1L: *M* = .013 [*d* = .64] | | .449 |
| | | | | Step-1R: *M* = .002 [*d* = .06] | | .498 |
| | | | | Step-2R: *M* = -.022 [*d* = -.92] | | .446 |
| | | | | Step-3R: *M* = .059 [*d* = 1.64] | | .245 |
| | | | | **GammaNS, mean difference [effect size]** | | |
| | | | | * *EL Hi-ADOS – EL Lo-ADOS* | | |
| | | | | Step-3L: *M* = -.025 [*d* = -1.19] | | .449 |
| | | | | Step-2L: *M* = -.001 [*d* = -.07] | | .492 |
| | | | | Step-1L: *M* = -.002 [*d* = -.09] | | .492 |
| | | | | Step-1R: *M* = -.026 [*d* = -1.04] | | .481 |
| | | | | Step-2R: *M* = .007 [*d* = .22] | | .492 |
| | | | | Step-3R: *M* = .048 [*d* = 1.25] | | .449 |
| | | | | * *EL Hi-ADOS – LL* | | |
| | | | | Step-3L: *M* = -.028 [*d* = -1.26] | | .449 |
| | | | | Step-2L: *M* = -.011 [*d* = -.53] | | .492 |
| | | | | Step-1L: *M* = -.003 [*d* = -.16] | | .492 |
| | | | | Step-1R: *M* = -.001 [*d* = -.03] | | .493 |
| | | | | Step-2R: *M* = -.018 [*d* = -.56] | | .492 |
| | | | | Step-3R: *M* = .060 [*d* = 1.48] | | .449 |
| | | | | * *EL Lo-ADOS – LL* | | |
| | | | | Step-3L: *M* = -.003 [*d* = -.16] | | .492 |
| | | | | Step-2L: *M* = -.009 [*d* = -.60] | | .492 |
| | | | | Step-1L: *M* = -.001 [*d* = -.10] | | .492 |
| | | | | Step-1R: *M* = .026 [*d* = 1.23] | | .449 |
| | | | | Step-2R: *M* = -.024 [*d* = -.98] | | .491 |
| | | | | Step-3R: *M* = .012 [*d* = .39] | | .492 |
| | | | | **GammaS, mean difference [effect size]** | | |
| | | | | * *EL Hi-ADOS – EL Lo-ADOS* | | |

| | | | | | | |
|---|---|---|---|---|---|---|
| | | | | Step-3L: *M* = -.025 [*d* = -1.21] | | .449 |
| | | | | Step-2L: *M* = -.013 [*d* = -.76] | | .492 |
| | | | | Step-1L: *M* = -.009 [*d* = -.52] | | .492 |
| | | | | Step-1R: *M* = -.044 [*d* = -1.83] | | .326 |
| | | | | Step-2R: *M* = .024 [*d* = .90] | | .492 |
| | | | | Step-3R: *M* = .067 [*d* = 1.97] | | .326 |
| | | | | * *EL Hi-ADOS – LL* | | |
| | | | | Step-3L: *M* = -.018 [*d* = -.82] | | .492 |
| | | | | Step-2L: *M* = -.016 [*d* = -.87] | | .492 |
| | | | | Step-1L: *M* = -.006 [*d* = -.31] | | .492 |
| | | | | Step-1R: *M* = -.039 [*d* = -1.55] | | .449 |
| | | | | Step-2R: *M* = .010 [*d* = .35] | | .492 |
| | | | | Step-3R: *M* = .069 [*d* = 1.90] | | .326 |
| | | | | * *EL Lo-ADOS – LL* | | |
| | | | | Step-3L: *M* = .007 [*d* = .42] | | .492 |
| | | | | Step-2L: *M* = -.003 [*d* = -.20] | | .492 |
| | | | | Step-1L: *M* = .003 [*d* = .23] | | .492 |
| | | | | Step-1R: *M* = .005 [*d* = .25] | | .492 |
| | | | | Step-2R: *M* = -.014 [*d* = -.65] | | .492 |
| | | | | Step-3R: *M* = .002 [*d* = .06] | | .492 |
| | | | | **AlphaNS, mean difference [effect size]** | | |
| | | | | * *EL Hi-ADOS – EL Lo-ADOS* | | |
| | | | | Step-3L: *M* = .013 [*d* = .61] | | .478 |
| | | | | Step-2L: *M* = .001 [*d* = .05] | | .478 |
| | | | | Step-1L: *M* = -.010 [*d* = -.41] | | .478 |
| | | | | Step-1R: *M* = -.052 [*d* = -1.61] | | .294 |
| | | | | Step-2R: *M* = -.005 [*d* = -.19] | | .478 |
| | | | | Step-3R: *M* = .052 [*d* = 1.26] | | .374 |
| | | | | * *EL Hi-ADOS – LL* | | |
| | | | | Step-3L: *M* = -.011 [*d* = -.49] | | .478 |
| | | | | Step-2L: *M* = -.002 [*d* = -.07] | | .478 |
| | | | | Step-1L: *M* = -.006 [*d* = -.53] | | .478 |
| | | | | Step-1R: *M* = -.013 [*d* = -.93] | | .478 |
| | | | | Step-2R: *M* = -.052 [*d* = -1.72] | | .294 |
| | | | | Step-3R: *M* = .110 [*d* = 2.49] | | .248 |
| | | | | * *EL Lo-ADOS – LL* | | |
| | | | | Step-3L: *M* = -.025 [*d* = -1.39] | | .327 |
| | | | | Step-2L: *M* = -.003 [*d* = -.16] | | .478 |
| | | | | Step-1L: *M* = -.003 [*d* = -.18] | | .478 |
| | | | | Step-1R: *M* = .020 [*d* = .77] | | .478 |
| | | | | Step-2R: *M* = -.046 [*d* = -1.98] | | .284 |
| | | | | Step-3R: *M* = .057 [*d* = 1.67] | | .294 |
| | | | | **AlphaS, mean difference [effect size]** | | |
| | | | | * *EL Hi-ADOS – EL Lo-ADOS* | | |
| | | | | Step-3L: *M* = .033 [*d* = 1.57] | | .294 |
| | | | | Step-2L: *M* = -.001 [*d* = -.07] | | .478 |
| | | | | Step-1L: *M* = .007 [*d* = .30] | | .478 |
| | | | | Step-1R: *M* = .003 [*d* = .07] | | .478 |
| | | | | Step-2R: *M* = -.065 [*d* = -1.99] | | .284 |
| | | | | Step-3R: *M* = .023 [*d* = .49] | | .478 |
| | | | | * *EL Hi-ADOS – LL* | | |
| | | | | Step-3L: *M* = .020 [*d* = .88] | | .478 |
| | | | | Step-2L: *M* = -.003 [*d* = -.14] | | .478 |
| | | | | Step-1L: *M* = -.003 [*d* = -.10] | | .478 |
| | | | | Step-1R: *M* = -.016 [*d* = -.38] | | .478 |
| | | | | Step-2R: *M* = -.026 [*d* = -.76] | | .478 |
| | | | | Step-3R: *M* = .027 [*d* = .56] | | .478 |
| | | | | * *EL Lo-ADOS – LL* | | |
| | | | | Step-3L: *M* = -.014 [*d* = -.78] | | .478 |

| | | | | | | |
|---|---|---|---|---|---|---|
| | | | | Step-2L: $M$ = -.002 [$d$ = -.09]<br>Step-1L: $M$ = -.010 [$d$ = -.50]<br>Step-1R: $M$ = -.018 [$d$ = -.58]<br>Step-2R: $M$ = .039 [$d$ = 1.45]<br>Step-3R: $M$ = .005 [$d$ = .12] | | .478<br>.478<br>.478<br>.327<br>.478 |
| Main result | **Results:** *Linking laterality of visual cortical functional connectivity and lateral global motion processing during infancy to later autism* | Table 1 | Main-effect model of ADOS2 ~ dbWPLI ["Model 0"] (OLS regression)<br><br>Main-effect model of ADOS2 ~ dbWPLI + GMLS ["Model 1"] (OLS regression)<br><br>Comparison of "Model 0" vs "Model 1" (LR test) | **<u>Regression coefficient [Effect size]</u>**<br>$\beta$(sex_Male) = .59 [part-□$^2$ = .043]<br>$\beta$(age) = -.02 [part-□$^2$ = .084]<br>$\beta$(status_LL) = .52 [part-□$^2$ = .025]<br>$\beta$(F_ThetaNS) = .56 [part-□$^2$ = .112]<br>$\beta$(F_GammaNS) = -.59 [part-□$^2$ = .107]<br>$\beta$(F_GammaS) = .39 [part-□$^2$ = .055]<br>$R^2$ = .322<br><br>$\beta$(sex_Male) = .40 [part-□$^2$ = .022]<br>$\beta$(age) = -.02 [part-□$^2$ = .089]<br>$\beta$(status_LL) = .34 [part-□$^2$ = .012]<br>$\beta$(F_ThetaNS) = .63 [part-□$^2$ = .152]<br>$\beta$(F_GammaNS) = -.57 [part-□$^2$ = .113]<br>$\beta$(F_GammaS) = .20 [part-□$^2$ = .015]<br>$\beta$(GMLS) = .47 [part-□$^2$ = .110]<br>$R^2$ = .397<br><br>**<u>npar \| LogLik \| Δdf \| $\chi^2$</u>**<br>Model 0: 6 -73.27<br>Model 1: 7 -70.77 1 5.014 | 43 | <br>.210<br>.077<br>.341<br>.039<br>.045<br>.155<br><br><br>.381<br>.072<br>.519<br>.017<br>.042<br>.474<br>.045<br><br><br><br><br>.025 |
| Main result | **Results:** *Linking laterality of visual cortical functional connectivity and lateral global motion processing during infancy to later autism* | Fig. 4a | Bivariate correlation (Pearson's $r$) | No point removed: $r$(F_GammaS x GMLS) = .30<br>Influential point removed: $r$(F_GammaS x ADOS2) = .42 | 49<br><br>48 | .038<br><br>.003 |
| Main result | **Results:** *Linking laterality of visual cortical functional connectivity and lateral global motion processing during infancy to later autism* | Fig. 4a | Main-effect model of GMLS ~ dbWPLI (OLS regression) | **<u>Regression coefficient [Effect size]</u>**<br>$\beta$(F_GammaS) = .36 [part-□$^2$ = .118]<br>$\beta$(F_ThetaS) = -.15 [part-□$^2$ = .025]<br>$\beta$(F_ThetaNS) = -.12 [part-□$^2$ = .012]<br>$\beta$(N_GammaNS) = .13 [part-□$^2$ = .015]<br>$\beta$(N_AlphaS) = -.14 [part-□$^2$ = .019]<br>$R^2$ = .161 | 49 | <br>.021<br>.301<br>.475<br>.423<br>.365 |
| Main result | **Results:** *Functional connectivity differences between stimulus types and between left and right hemispheres* | Figs. 4b-c<br>Ext. Data Figs. 3c-d | Across-stimulus difference test of flipped, norm. AoI connectivity (Permutation test + FDR correct.)<br>*<u>Notes:</u>*<br>[i])EL-High ADOS<br>[ii])EL-Low ADOS | **<u>Theta, mean difference [effect size]</u>**<br>* *Social – NonSocial; EL-High ADOS*<br>Step-3L: $M$ = .006 [$d$ = .22]<br>Step-2L: $M$ = .006 [$d$ = .28]<br>Step-1L: $M$ = .026 [$d$ = .99]<br>Step-1R: $M$ = .050 [$d$ = 1.31]<br>Step-2R: $M$ = .063 [$d$ = 2.09]<br>Step-3R: $M$ = -.151 [$d$ = -3.04]<br>* *Social – NonSocial; EL-Low ADOS* | 59<br>11[i)]<br>28[ii)]<br>20[iii)] | <br><br>.481<br>.148<br>.352<br>.148<br>.210<br>.002 |

| | | | | | | |
|---|---|---|---|---|---|---|
| | | | iii)LL | Step-3L: *M* = -.019 [*d* = -1.37] | | .210 |
| | | | | Step-2L: *M* = -.047 [*d* = -4.22] | | .210 |
| | | | | Step-1L: *M* = .010 [*d* = .74] | | .485 |
| | | | | Step-1R: *M* = -.028 [*d* = -1.49] | | .352 |
| | | | | Step-2R: *M* = -.010 [*d* = -.67] | | .148 |
| | | | | Step-3R: *M* = .094 [*d* = 3.83] | | .017 |
| | | | | ** Social – NonSocial; LL* | | |
| | | | | Step-3L: *M* = .021 [*d* = 1.17] | | .229 |
| | | | | Step-2L: *M* = -.033 [*d* = -2.22] | | .485 |
| | | | | Step-1L: *M* = -.002 [*d* = -.13] | | .352 |
| | | | | Step-1R: *M* = -.037 [*d* = -1.44] | | .352 |
| | | | | Step-2R: *M* = .033 [*d* = 1.66] | | .352 |
| | | | | Step-3R: *M* = .019 [*d* = .57] | | .485 |
| | | | | **Gamma, mean difference [effect size]** | | |
| | | | | ** Social – NonSocial; EL-High ADOS* | | |
| | | | | Step-3L: *M* = .001 [*d* = .06] | | .462 |
| | | | | Step-2L: *M* = .002 [*d* = .16] | | .462 |
| | | | | Step-1L: *M* = -.017 [*d* = -1.04] | | .462 |
| | | | | Step-1R: *M* = -.043 [*d* = -2.40] | | .462 |
| | | | | Step-2R: *M* = .034 [*d* = 2.03] | | .462 |
| | | | | Step-3R: *M* = .023 [*d* = .93] | | .462 |
| | | | | ** Social – NonSocial; EL-Low ADOS* | | |
| | | | | Step-3L: *M* = .001 [*d* = .11] | | .462 |
| | | | | Step-2L: *M* = .014 [*d* = 1.98] | | .462 |
| | | | | Step-1L: *M* = -.010 [*d* = -1.16] | | .462 |
| | | | | Step-1R: *M* = -.025 [*d* = -2.83] | | .462 |
| | | | | Step-2R: *M* = .016 [*d* = 1.94] | | .495 |
| | | | | Step-3R: *M* = .004 [*d* = .34] | | .462 |
| | | | | ** Social – NonSocial; LL* | | |
| | | | | Step-3L: *M* = -.009 [*d* = -1.09] | | .462 |
| | | | | Step-2L: *M* = .007 [*d* = .76] | | .462 |
| | | | | Step-1L: *M* = -.014 [*d* = -1.31] | | .462 |
| | | | | Step-1R: *M* = -.004 [*d* = -.37] | | .462 |
| | | | | Step-2R: *M* = .006 [*d* = .57] | | .462 |
| | | | | Step-3R: *M* = .015 [*d* = .90] | | .462 |
| | | | | **Alpha, mean difference [effect size]** | | |
| | | | | ** Social – NonSocial; EL-High ADOS* | | |
| | | | | Step-3L: *M* = -.010 [*d* = -.48] | | .458 |
| | | | | Step-2L: *M* = -.028 [*d* = -1.15] | | .497 |
| | | | | Step-1L: *M* = .026 [*d* = 1.00] | | .474 |
| | | | | Step-1R: *M* = .060 [*d* = 1.78] | | .458 |
| | | | | Step-2R: *M* = -.021 [*d* = -.71] | | .467 |
| | | | | Step-3R: *M* = -.028 [*d* = -.66] | | .458 |
| | | | | ** Social – NonSocial; EL-Low ADOS* | | |
| | | | | Step-3L: *M* = -.030 [*d* = -2.82] | | .497 |
| | | | | Step-2L: *M* = -.025 [*d* = -2.02] | | .497 |
| | | | | Step-1L: *M* = .009 [*d* = .70] | | .474 |
| | | | | Step-1R: *M* = .006 [*d* = .36] | | .390 |
| | | | | Step-2R: *M* = .038 [*d* = 2.59] | | .063 |
| | | | | Step-3R: *M* = .002 [*d* = .07] | | .474 |
| | | | | ** Social – NonSocial; LL* | | |
| | | | | Step-3L: *M* = -.041 [*d* = -2.90] | | .467 |
| | | | | Step-2L: *M* = -.026 [*d* = -1.61] | | .497 |
| | | | | Step-1L: *M* = .016 [*d* = .90] | | .497 |
| | | | | Step-1R: *M* = .045 [*d* = 1.97] | | .467 |
| | | | | Step-2R: *M* = -.047 [*d* = -2.37] | | .074 |
| | | | | Step-3R: *M* = .054 [*d* = 1.88] | | .390 |

| | | | | | | |
|---|---|---|---|---|---|---|
| Main result | **Results:**<br>*Functional connectivity differences between stimulus types and between left and right hemispheres* | Ext. Data Figs. 3a-b | Across-stimulus difference test of absolute (raw) AoI connectivity (Permutation test + FDR correct.)<br>*Notes:*<br>[i)]EL-High ADOS<br>[ii)]EL-Low ADOS<br>[iii)]LL | **Theta, mean difference [effect size]**<br>* *Social – NonSocial; EL-High ADOS*<br>Step-3L: *M* = -.008 [*d* = -1.28]<br>Step-2L: *M* = -.002 [*d* = -.38]<br>Step-1L: *M* = .004 [*d* = .92]<br>Step-1R: *M* = .002 [*d* = .39]<br>Step-2R: *M* = .001 [*d* = .24]<br>Step-3R: *M* = -.004 [*d* = -.63]<br>* *Social – NonSocial; EL-Low ADOS*<br>Step-3L: *M* = .007 [*d* = 1.96]<br>Step-2L: *M* = .001 [*d* = .46]<br>Step-1L: *M* = .003 [*d* = 1.26]<br>Step-1R: *M* = .005 [*d* = 1.61]<br>Step-2R: *M* = .004 [*d* = 1.22]<br>Step-3R: *M* = .005 [*d* = 1.41]<br>* *Social – NonSocial; LL*<br>Step-3L: *M* = -.002 [*d* = -.47]<br>Step-2L: *M* = -.004 [*d* = -1.02]<br>Step-1L: *M* = .000 [*d* = .15]<br>Step-1R: *M* = -.004 [*d* = -.84]<br>Step-2R: *M* = -.004 [*d* = -.89]<br>Step-3R: *M* = .001 [*d* = .20]<br><br>**Gamma, mean difference [effect size]**<br>* *Social – NonSocial; EL-High ADOS*<br>Step-3L: *M* = .002 [*d* = .36]<br>Step-2L: *M* = .002 [*d* = .47]<br>Step-1L: *M* = -.002 [*d* = -.39]<br>Step-1R: *M* = .000 [*d* = -.13]<br>Step-2R: *M* = -.001 [*d* = -.33]<br>Step-3R: *M* = .001 [*d* = .28]<br>* *Social – NonSocial; EL-Low ADOS*<br>Step-3L: *M* = .007 [*d* = 2.41]<br>Step-2L: *M* = .008 [*d* = 2.95]<br>Step-1L: *M* = .003 [*d* = 1.43]<br>Step-1R: *M* = .005 [*d* = 2.33]<br>Step-2R: *M* = .003 [*d* = 1.43]<br>Step-3R: *M* = .004 [*d* = 2.69]<br>* *Social – NonSocial; LL*<br>Step-3L: *M* = .003 [*d* = .88]<br>Step-2L: *M* = .004 [*d* = 1.04]<br>Step-1L: *M* = -.001 [*d* = -.23]<br>Step-1R: *M* = .003 [*d* = 1.15]<br>Step-2R: *M* = .002 [*d* = .93]<br>Step-3R: *M* = .002 [*d* = .98]<br><br>**Alpha, mean difference [effect size]**<br>* *Social – NonSocial; EL-High ADOS*<br>Step-3L: *M* = -.002 [*d* = -.83]<br>Step-2L: *M* = -.004 [*d* = -1.21]<br>Step-1L: *M* = .001 [*d* = .25]<br>Step-1R: *M* = .003 [*d* = .85]<br>Step-2R: *M* = -.002 [*d* = -.52]<br>Step-3R: *M* = -.002 [*d* = -.76]<br>* *Social – NonSocial; EL-Low ADOS*<br>Step-3L: *M* = -.005 [*d* = -2.86]<br>Step-2L: *M* = -.004 [*d* = -2.19]<br>Step-1L: *M* = -.006 [*d* = -2.50]<br>Step-1R: *M* = -.004 [*d* = -1.56]<br>Step-2R: *M* = -.001 [*d* = -.35]<br>Step-3R: *M* = .002 [*d* = 1.35] | 59<br>11[i)]<br>28[ii)]<br>20[iii)] | .<br><br>.373<br>.467<br>.467<br>.467<br>.467<br>.373<br><br>.373<br>.373<br>.467<br>.373<br>.373<br>.373<br><br>.437<br>.437<br>.437<br>.373<br>.373<br>.467<br><br><br><br>.473<br>.473<br>.473<br>.473<br>.473<br>.473<br><br>.473<br>.473<br>.473<br>.473<br>.473<br>.473<br><br>.473<br>.495<br>.473<br>.473<br>.473<br>.473<br><br><br><br>.350<br>.350<br>.350<br>.347<br>.350<br>.350<br><br>.473<br>.473<br>.473<br>.473<br>.473<br>.473 |

| | | | | | | |
|---|---|---|---|---|---|---|
| | | | | ** Social – NonSocial; LL*<br>Step-3L: *M* = -.003 [*d* = -1.21]<br>Step-2L: *M* = -.009 [*d* = -3.23]<br>Step-1L: *M* = -.003 [*d* = -.90]<br>Step-1R: *M* = -.004 [*d* = -1.21]<br>Step-2R: *M* = -.009 [*d* = -3.04]<br>Step-3R: *M* = -.003 [*d* = -1.19] | | <br>.350<br>.350<br>.444<br>.350<br>.347<br>.350 |
| Main result | **Results:** *Functional connectivity differences between stimulus types and between left and right hemispheres* | Fig. 4d | Left-/right-sidedness difference test of dbWPLI variables (Wilcoxon test + FDR correction)<br>*Notes:*<br>i)EL-High ADOS<br>ii)EL-Low ADOS<br>iii)LL | **Theta, mean difference [effect size]**<br>** Right – Left; NonSocial stimulus*<br>EL-HiADOS: *M* = -.024 [*d* = -.11]<br>EL-LoADOS: *M* = .050 [*d* = .47]<br>LL: *M* = -.021 [*d* = - .23]<br>** Right – Left; Social stimulus*<br>EL-HiADOS: *M* = -.042 [*d* = -.39]<br>EL-LoADOS: *M* = -.033 [*d* = -.24]<br>LL: *M* = -.022 [*d* = -.22]<br><br>**Gamma, mean difference [effect size]**<br>** Right – Left; NonSocial stimulus*<br>EL-HiADOS: *M* = -.037 [*d* = -.27]<br>EL-LoADOS: *M* = -.094 [*d* = -1.03]<br>LL: *M* = -.028 [*d* = -.32]<br>** Right – Left; Social stimulus*<br>EL-HiADOS: *M* = -.053 [*d* = -.38]<br>EL-LoADOS: *M* = -.067 [*d* = -.82]<br>LL: *M* = -.074 [*d* = -.82]<br><br>**Alpha, mean difference [effect size]**<br>** Right – Left; NonSocial stimulus*<br>EL-HiADOS: *M* = -.079 [*d* = -.68]<br>EL-LoADOS: *M* = -.053 [*d* = -.50]<br>LL: *M* = -.018 [*d* = -.23]<br>** Right – Left; Social stimulus*<br>EL-HiADOS: *M* = -.039 [*d* = -.38]<br>EL-LoADOS: *M* = .047 [*d* = .40]<br>LL: *M* = -.011 [*d* = -.08] | 59<br>11i)<br>28ii)<br>20iii) | <br><br>.623<br>.131<br>.623<br><br>.616<br>.616<br>.677<br><br><br><br>.897<br>.004<br>.616<br><br>.616<br>.031<br>.131<br><br><br><br>.419<br>.616<br>.616<br><br>.616<br>.616<br>.989 |
| Supplementary result | **Results:** *Linking laterality of visual cortical functional connectivity and lateral global motion processing during infancy to later autism* | Last paragraph of section | Main-effect model of ADOS2 ~ GLMS (OLS regression) | **Regression coefficient [Effect size]**<br>$\beta$(sex_Male) = 1.17 [part-$\eta^2$ = .101]<br>$\beta$(age) = .00 [part-$\eta^2$ = .000]<br>$\beta$(status_LL) = -1.09 [part-$\eta^2$ = .065]<br>$\beta$(GMLS) = .53 [part-$\eta^2$ = .074]<br>$\beta$(GFLS) = .25 [part-$\eta^2$ = .019]<br>$\beta$(LMLS) = .02 [part-$\eta^2$ = .000]<br>$\beta$(LFLS) = -.30 [part-$\eta^2$ = .021]<br>$R^2$ = .233 | 82 | <br>.005<br>.898<br>.026<br>.018<br>.236<br>.914<br>.210 |

***Abbreviations:*** FC_ = Far-Connectivity; NC_ = Near-Connectivity; [Band]NS = NonSocial stimuli; [Band]S = Social stimuli; dbWPLI = debiased Weighted Phase Lag Index; GMLS = Global Motion Laterality Score; EL = Elevated Likelihood for ASD; LL = Low Likelihood for ASD; LR = Likelihood Ratio; npar = no. of model parameters; LogLik = Log-Likelihood; df = degrees of freedom; GFLS = Global Form Laterality Score; LMLS = Local Motion Laterality Score; LFLS = Local Form Laterality Score

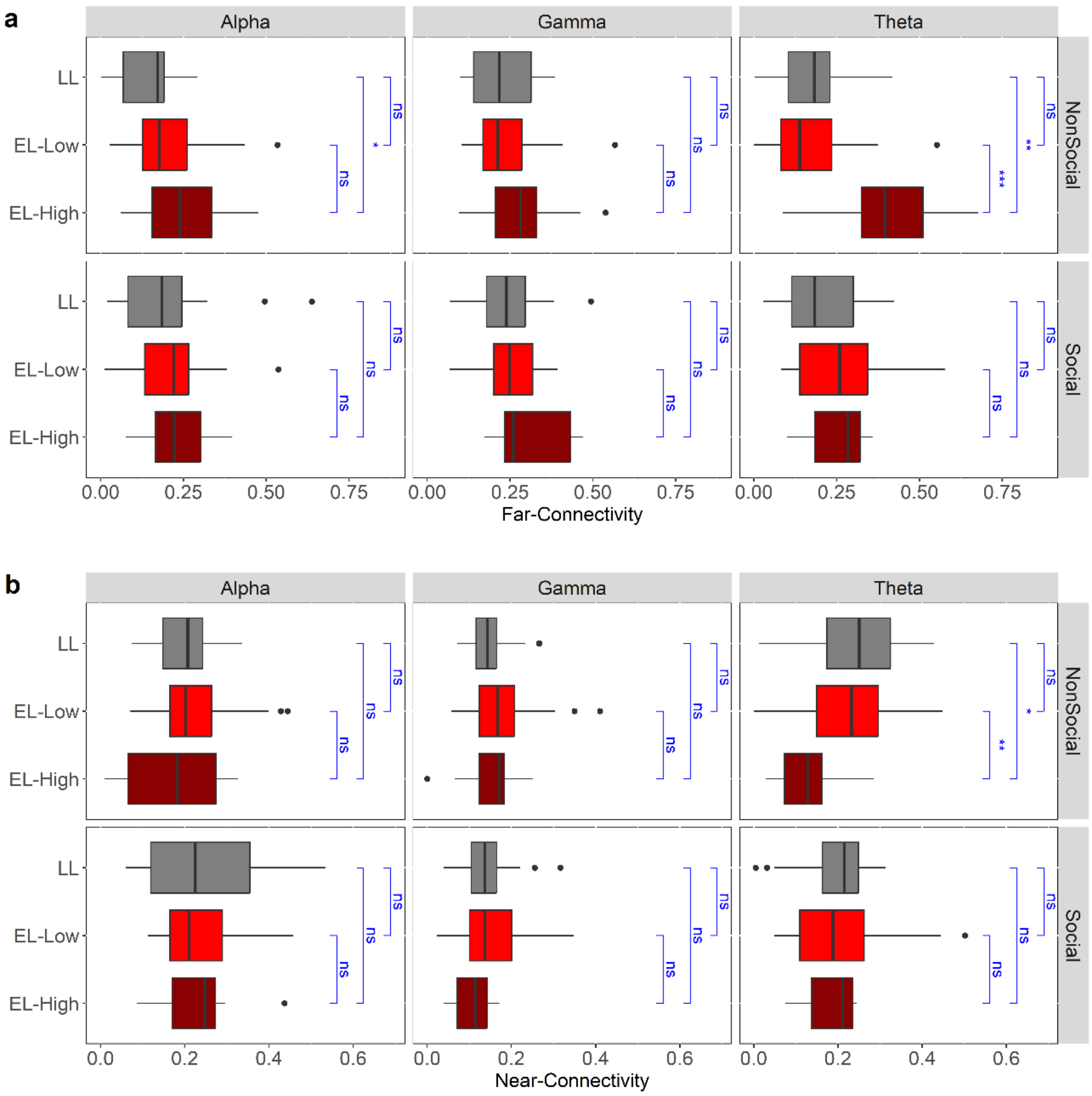


**Supplementary Figure 1 | Group analysis of dbWPLI Near & Far Connectivity variables in six experimental conditions. a.** Group differences in Far Connectivity variables in the six experimental conditions. Significant differences were found in Theta-NonSocial condition between the EL-High ADOS group and the other two groups, but not between these two later groups, and in Alpha Non-Social condition between the two EL groups. No correction was done on the *p*-values here. FDR corrections of *p*-values resulted in the elimination of significance in Alpha-NonSocial but not in Theta-NonSocial conditions (see also **Fig. 3b**). **b.** Group differences in Near Connectivity variables in the six experimental conditions. Significant differences were found in Theta-NonSocial condition between the EL-High ADOS group and the other two groups, but not between these two later groups. Again, no *p*-value correction was done here. FDR corrections of *p*-values resulted in no significant difference in any conditions. See **Supplementary Table 5** for all numerical details. Boxplots show the sample median, and the first and third quartiles; whiskers show minimum and maximum (± 1.5 s.d.); dots are outliers (± 1.96 s.d.). (Notes: EL = Elevated Likelihood; LL = Low Likelihood; EL-High = EL-High ADOS; EL-Low = EL-Low ADOS; ns = $p > .05$; * = $p < .05$; ** = $p < .01$; *** = $p < .001$).

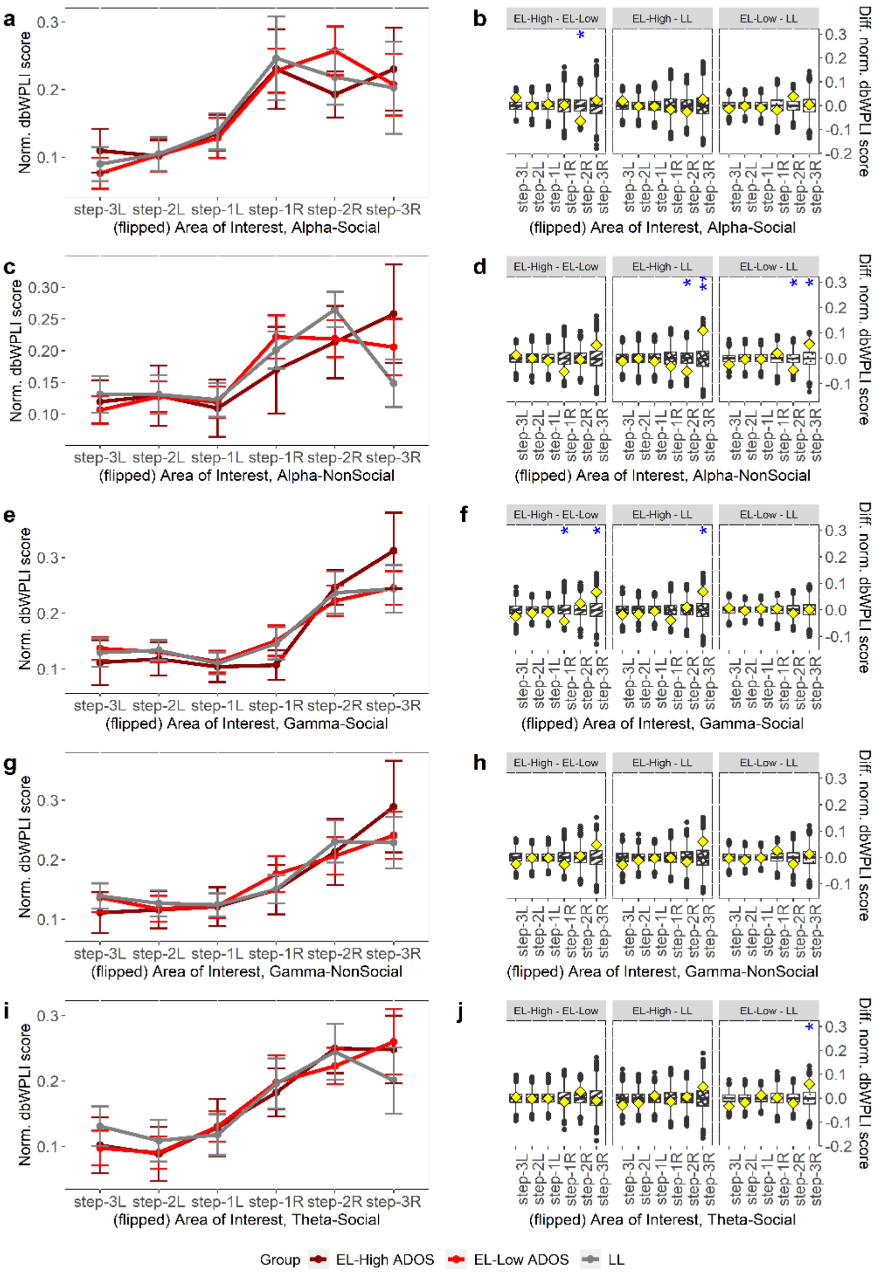


**Supplementary Figure 2 | Permutation tests of across-group differences in five experimental conditions based on the flipped topographies.** Similar to **Figs. 3c-d**, but for **a-b.** Alpha-Social condition, showing a difference in step-2R AoI between the EL groups; **c-d.** Alpha-NonSocial condition, showing a difference in step-2R and step-3R AoIs between the each EL group and LL group, but not between the two EL groups; **e-f.** Gamma-Social condition, showing a difference in step-1R and step-3R AoIs between the two EL groups, and in step-3R AoI between the EL-High ADOS and LL groups; **g-h.** Gamma-NonSocial condition, showing no difference; **i-j.** Theta-Social condition, showing a difference in step-3R AoI between the EL-Low ADOS and LL groups. No *p*-value correction was done. FDR correction for multiple testing resulted in no significance found in these 5 conditions (also see **Figs. 3c-d**). Complete numerical details in **Supplementary Table 5**. (*Note*: boxplots are simulated distributions using 10,000 randomization of the group labels, diamonds actual point values in our data, stars mark significance).

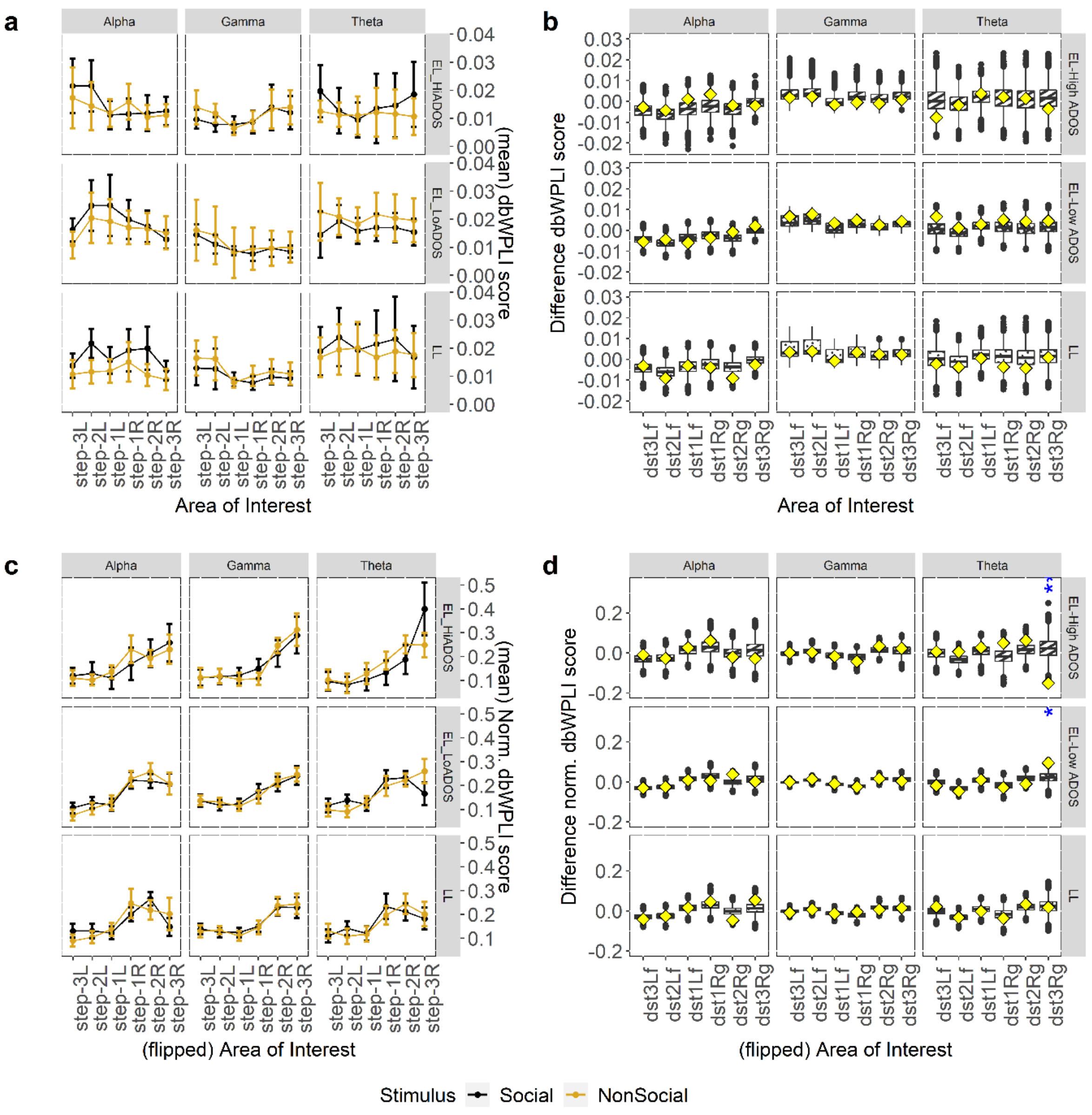


**Supplementary Figure 3 | Across-stimulus topographic connectivity differences in each frequency band and group. a.** Plots of average raw (unnormalized, unflipped) topographic dbWPLI scores across the two stimuli for each group and frequency band. **b.** Permutation tests of across-stimulus differences for the topographies in **a**, showing no significant difference (all $p > .300$) for any group or band (boxplots are simulated distributions using 10,000 randomization of the group labels, diamonds actual point values in our data, stars mark significance). **c-d.** Similar to **a-b**, but for the flipped normalized dbWPLI scores, where a significant difference in 3-step AoI in theta band were revealed for both EL groups (High ADOS: $d = -3.05$, $^{**}p = .002$, $n = 11$; Low ADOS: $d = 3.82$, $^{*}p = .017$, $n = 28$) by the permutation tests. All $p$-values were FDR-corrected for multiple testing. (Notes: EL = Elevated Likelihood; LL = Low Likelihood; step-$d$L/-R = step-$d$Left/-Right, $d = 1, 2, 3$; * = $p < .05$; ** = $p < .01$; *** = $p < .001$). Complete numerical details in **Supplementary Table 5**.

### References

1 Hardiansyah, I. *et al.* Global motion processing in infants' visual cortex and the emergence of autism. *Communications Biology* **6**, 1-10 (2023).

2 Mullen, E. M. *Mullen scales of early learning*. (AGS Circle Pines, MN, 1995).
